# Tissue and cellular spatiotemporal dynamics in colon aging

**DOI:** 10.1101/2024.04.22.590125

**Authors:** Aidan C. Daly, Francesco Cambuli, Tarmo Äijö, Britta Lötstedt, Nemanja Marjanovic, Olena Kuksenko, Matthew Smith-Erb, Sara Fernandez, Daniel Domovic, Nicholas Van Wittenberghe, Eugene Drokhlyansky, Gabriel K Griffin, Hemali Phatnani, Richard Bonneau, Aviv Regev, Sanja Vickovic

**Affiliations:** New York Genome Center, New York, NY, USA; Center for Computational Biology, Flatiron Institute, New York, NY, USA; Klarman Cell Observatory Broad Institute of MIT and Harvard, Cambridge, MA, USA; Science for Life Laboratory, Department of Gene Technology, KTH Royal Institute of Technology, Stockholm, Sweden; Department of Biology, Massachusetts Institute of Technology, Cambridge, MA, USA; Department of Neurology, Columbia University Irving Medical Center, New York, NY, USA; Department of Pathology, Brigham and Women’s Hospital, Boston, MA, USA; Center for Data Science, New York University, New York, NY, USA; Genentech, 1 DNA Way, South San Francisco, CA, USA; Department of Biomedical Engineering and Herbert Irving Institute for Cancer Dynamics, Columbia University, New York, NY, USA; Science for Life Laboratory, Department of Immunology, Genetics and Pathology, Beijer Laboratory for Gene and Neuro Research, Uppsala University, Uppsala, Sweden

## Abstract

Tissue structure and molecular circuitry in the colon can be profoundly impacted by systemic age-related effects, but many of the underlying molecular cues remain unclear. Here, we built a cellular and spatial atlas of the colon across three anatomical regions and 11 age groups, encompassing ∼1,500 mouse gut tissues profiled by spatial transcriptomics and ∼400,000 single nucleus RNA-seq profiles. We developed a new computational framework, cSplotch, which learns a hierarchical Bayesian model of spatially resolved cellular expression associated with age, tissue region, and sex, by leveraging histological features to share information across tissue samples and data modalities. Using this model, we identified cellular and molecular gradients along the adult colonic tract and across the main crypt axis, and multicellular programs associated with aging in the large intestine. Our multi-modal framework for the investigation of cell and tissue organization can aid in the understanding of cellular roles in tissue-level pathology.

## INTRODUCTION

A typical colon extends >12 cm in mice and >1.5 meters in humans^1,2^, with considerable variance in length, thickness, and folding, impacted by multiple variables, including age, sex, weight, and diet. The inner lumen of the large intestine is punctuated by millions of invaginations, each harboring a colonic crypt, as the key anatomic unit responsible for its continuous regeneration and differentiation^3^. Underlying the mucosal epithelium, the submucosa hosts lymphoid clusters, nerve fibers, and the lymphovasculature, while the outer muscular wall enables peristaltic motility. From the ceacum to the rectum, the colon carries out spatially confined regional functions, which emerge postnatally and are required for the digestion of solid food^4,5^. During aging, the decline of colonic function is accompanied by dysbiosis and excessive epithelial permeability, allowing the gut microbiota to infiltrate the lumen^6^ and causing a generalized and protracted inflammatory state^7^, and the emergence of common pathologies, including constipation, diverticulitis, malnutrition and colorectal cancer^8^.

Despite the crucial importance of colon function, the cellular and molecular features associated with functional diversity across colonic regions, the crypt axis, and major lifespan stages have not yet been comprehensively characterized. In recent years, single cell and single nucleus profiling of the mouse and human intestines^9–20^ has discovered and classified cell types and functions in the gut during development^15,20^, in adults^16,17,19^, and in aging^18,21^, but with limited spatial context. Spatial *in situ* profiling methods^22–33^ are poised to address this gap, through either targeted or genome-wide profiling. However, robust computational frameworks for spatial analysis of large tissue cohorts are still lacking. For example, many spatial analysis methods reduce noise by smoothing gene expression data across neighboring spots or cellular neighborhoods, at the risk of true signal loss^34–36^. Most methods for testing spatial differential expression through clustering^37,38^ or explicit modeling^39,40^ are applied only to single tissue sections, and those that integrate data from multiple tissue sections^41–44^ are usually limited to the alignment of serial sections from the same tissue. Other methods, focused on deconvolution of multi-cellular spatially resolved measurements to the single cell level (through Bayesian modeling^45–47^, non-negative matrix factorization (NMF)^48^, or deep learning^49^), typically use little or no information about tissue anatomy or histology, which may yield biologically unrealistic results and limit their efficacy^50,51^. Furthermore, while a common coordinate system^52^ can facilitate integration, it is more challenging in large tissues like the colon that lacks a strict stereotypical architecture. Thus, to characterize the molecular and cellular variation underlying functional variation in the colon, both spatial profiling of large tissue cohorts, and computational means to integrate these data across space and age are needed.

Here, we created a comprehensive experimental and computational framework to construct an integrated cell and tissue atlas of the mouse colon across temporal, anatomical, morphological variation, by combining Spatial Transcriptomics (ST)^31^ and single nucleus RNA-seq (snRNA-seq)^17^. We define the relative abundance of cell types using multi-modal estimation, address missing data imputation and technical noise correction through information sharing across tissue sections and use explicit, hierarchical modeling of spatial and covariate-specific effects on cellular gene expression for Bayesian hypothesis testing across cell types, tissue regions, ages, or other covariate groups informed by both snRNA-seq and ST data. Our work provides important insights into tissue and cell-level function and organization and serves as an important reference for understanding the biology of aging.

## RESULTS

### A spatiotemporal atlas of the colon

To build a comprehensive atlas, we collected colonic specimens from proximal to distal anatomical regions through three major phases of the mammalian lifespan: juvenile (<4 weeks of age in the mouse), adulthood (6 –12 weeks) and aging (6 months – 2 years) (**Fig. 1a,b**) and profiled them by snRNA-seq and ST (**Fig. 1c,d**). The cellular branch of the atlas encompassed ∼400,000 snRNA-seq profiles from 21 specimens, which we partitioned and annotated at first into 17 major subsets of epithelial cells (intestinal stem cells (ISCs), transamplifying (TAs), cycling TAs, colonocytes, goblets, neuroendocrines and tufts), immune cells (B, T, macrophages), stromal cells (vascular, lymphatics, fibroblasts, trophocytes), enteric neurons, glia, and smooth muscle cells (SMC)^17,21,53^ **(Fig. 1f, Extended Data Fig. 1, Extended Data Table 1, Methods**). The spatial branch comprised ∼1,500 sections from 65 mice and ∼66,500 spatially barcoded spots, each quantifying the expression of 12,976 genes, sampling, on average, 24 tissue sections, 977 spots and 35,730 annotated cell segments from each mouse colon. These spanned 66 conditions across combinations of age, sex, colonic region and morphological regions of interest (MROIs) (**Fig. 1g-i, Extended Data Table 2**).

**Fig. 1.**
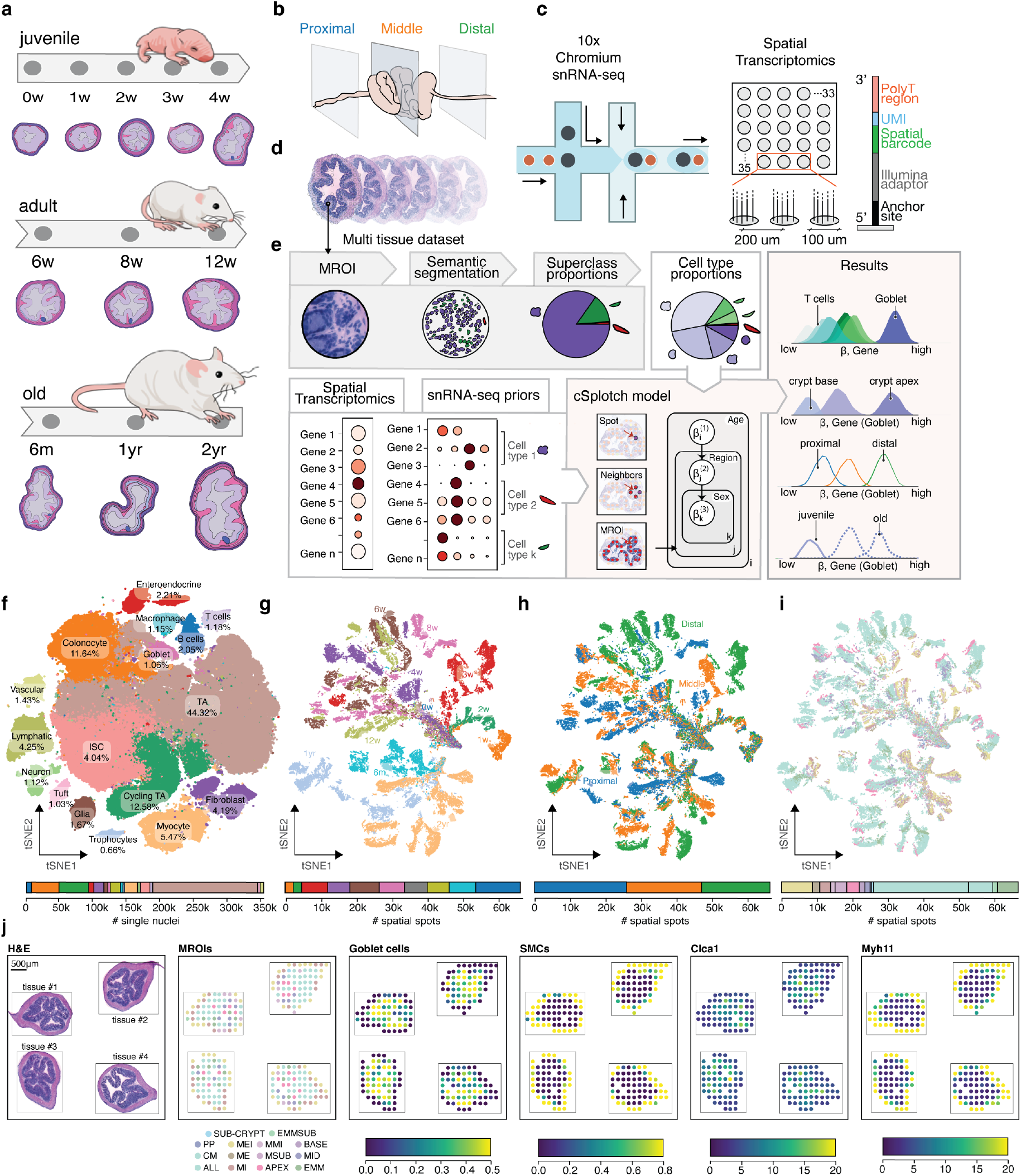
A cellular and spatial atlas of the mouse colon across regions and ages. **(a**-**e**) Study design overview. (**a**) Sampling time points: Birth/juvenile (0-4w), adulthood (6-12w) and aging (6m-2yr). (**b**) Sampling regions: proximal, middle, and distal colon. (**c)** Profiling methods: snRNA-seq (left) and barcoded spatial transcriptomics (right). (**d,e**) Analysis. Multi-tissue dataset (d) served as input to cSplotch (e), which uses histological (“MROI”, “Semantic segmentation”) and expression (“Spatial Transcriptomics”, “snRNA-seq priors”) data to estimate the abundance of each snRNA-seq cell type in each spatial spot, then employs hierarchical Bayesian modeling (“cSplotch model”) to infer cell type-specific gene expression conditioned on age, region, sex, and MROI annotation (“Results”). (**f**) Single nucleus atlas. t-Distributed Stochastic Neighbor Embedding (t-SNE) of 352,195 snRNA-seq profiles colored by cell type cluster (top), and their relative abundances (stacked bars, bottom). (**g-i**) Spatial atlas. tSNE embedding of 66,481 spatial transcriptomics spot profiles colored by age group (**g**), colon section (**h**), and MROI (**i**) (top), along with relative abundances (stacked bars, bottom). (**j**) Example sections. From left: Tissue sections (3w; proximal colon) processed with spatial transcriptomics stained with Hematoxilin and Eosin (H&E) (leftmost), and with measured spots colored by MROI annotation, goblet and smooth muscle cells (SMCs) inferred proportions, and inferred expression rates (*<u>λ</u>*) for specific genes, Scale bar: 500μm.

### cSplotch infers cell type compositions and gene expression rates from ST, histology and snRNA-seq

To accurately detect spatial gene expression changes across multiple tissue samples, individuals and conditions in our atlas, we developed cSplotch, a novel hierarchical Bayesian probabilistic model that uses both histological images and snRNA-seq to infer location- and covariate-dependent cell type-specific gene expression profiles from multicellular ST data (**Fig. 1e**). Overall, cSplotch consists of two major steps. First, it infers the cellular composition in each spatial spot from a multi-tissue dataset using morphological regions of interest (MROIs), cell level morphological data, and snRNA-seq profiles. Second, it uses these cell type compositions to infer MROI- and covariate-specific expression rates for each gene in each cell type; these rates can then be used to test for differential expression across location, condition, or cell type in an entire atlas or to infer multicellular programs (MCPs^54^) of expression patterns from multiple cell types coordinated across samples.

Specifically, to infer cell composition, we first annotated each spatial spot with an MROI label (**Methods**), segmented nuclei from the histology image, and annotated each nuclear segment with one of five morphological cell superclass labels (**Fig. 2a**), using a conditioned semantic segmentation workflow employing structural, anatomical and neighborhood features (**Extended Data Fig. 2, Extended Data Table 3, Methods**). Across held-out test sets for young (≤4 weeks old) and adult (≥6 weeks old) mice and all anatomical regions, we correctly assigned 85% of all nuclear areas when compared against pathologist superclass annotations (**Extended Data Fig. 2c-d**). Next, for each spot, cSplotch uses NMF^52,55^ to infer a combination of snRNA-seq profiles whose aggregate marker gene expression profile fits the observed expression measurement, but in a manner constrained by the morphological cell composition inferred from histology in the previous step (**Methods, Extended Data Table 4**). Without introducing any morphological cell composition constraints (*α* = 0), while gene expression profiles were well-reconstructed (**Extended Data Fig. 3a**, blue line), the morphological cell type proportions were not, resulting in large numbers of deconvolved transcriptional cell types assigned to each spot (**Extended Data Fig. 3b,c**, blue lines). Once constrained by the inferred morphological cell type proportions (*α*>0, **Extended Data Fig. 3c**, orange, green and red lines), the cell composition reconstruction improved without loss of expression reconstruction accuracy. cSplotch performed well in both decomposing the expression profiles and recovering cell compositions consistent with pathologist annotations and known features of the tissue. For example, cell composition estimates for 13 randomly selected spots from different MROIs from a tissue segment of the distal 8-week colon, agreed well with ground truth pathologist annotations and well known patterns of tissue organization in terms of cell type proportions (**Fig. 2a**). These included high SMC content in the *muscularis* regions, high goblet cell content in epithelial layers and high B cell content in the *Peyer’s patch* (PP) (**Fig. 2a**).

**Fig. 2.**
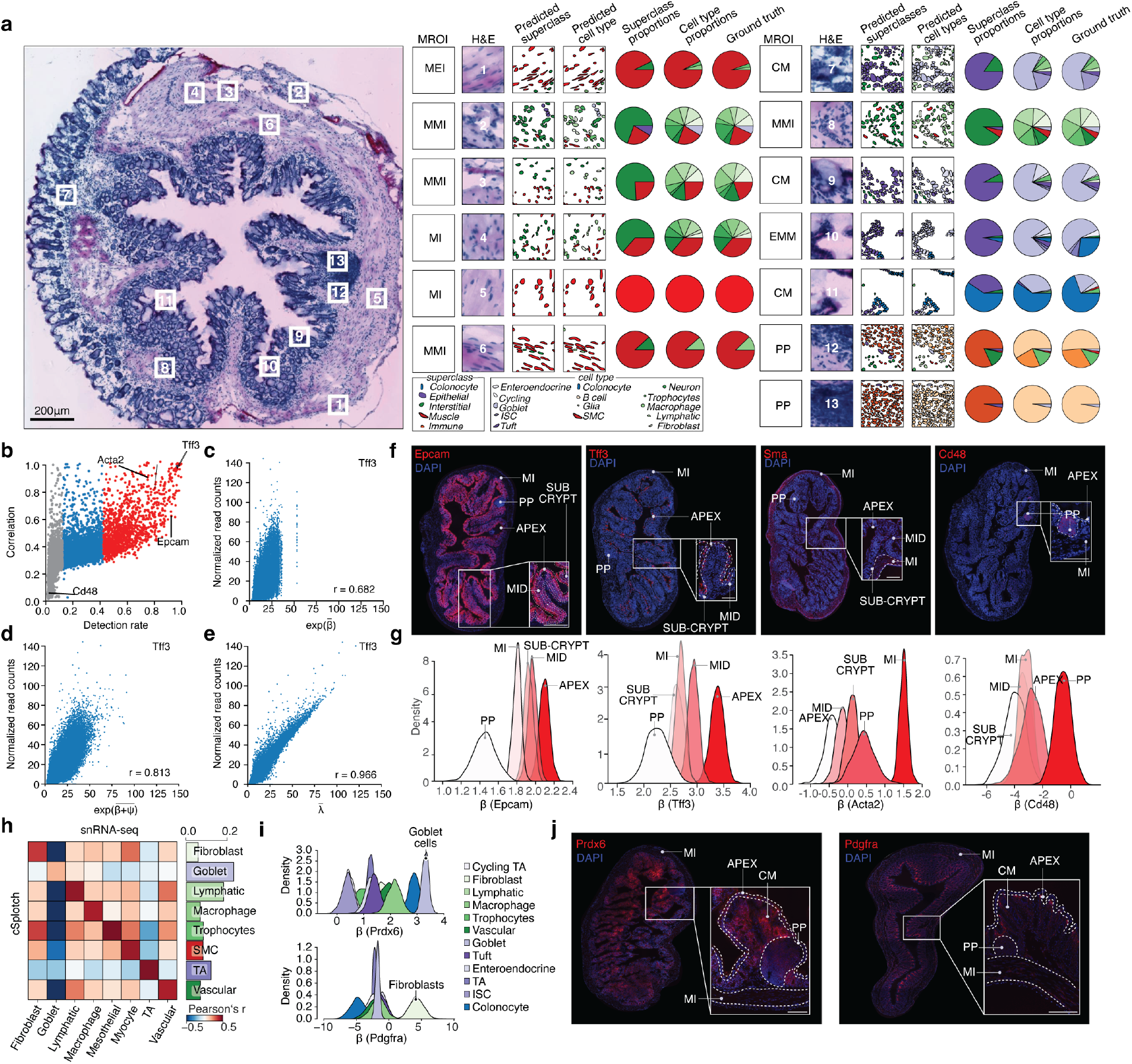
Deconvolutional modeling using histological and expression features. **(a**) Agreement of morphology-informed deconvolution with manual cell type annotation. Thirteen H&E patches from an example ST array (left; numbered boxes; scale bar: 200μm) with their (from left) MROI annotation, H&E image patch, semantic segmentation of morphological superclasses (color code; bottom left) and snRNA-seq cell types (color code; bottom right), and relative proportions of predicted superclasses (pie charts), predicted cell types, and manually labeled cell types (“ground truth”). (**b-e**) cSplotch validation. (**b**) Pearson correlation coefficients (y axis) between the expression rate estimated by cSplotch (*λ*) and TPM normalized values in ST for the same spots scattered against the detection rate in ST (percentage of spots with non-zero measurements, x axis) for each of 12,796 genes (dots). Red/grey: top 10% / bottom 50% in detection rate. (**c-e**) Variance in *Tff3* gene expression (y axis, normalized read counts) across spots (n=66,481, dots) explained by region alone (mean 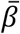, x axis, **c**); region and location (mean (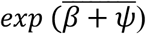, x axis, **d**), or all components (mean 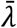, x axis, **e**) (all x axis values are. Lower right: Pearson’s *r*. (**f-g**) IF validation of spatial gene expression trends predicted by cSplotch. (**f,g**) Immunofluorescence (IF) validation. (**f**) IF of four gene products (*Epcam, Tff3, Sma*, and *Cd48*; also highlighted in (b)) in proximal colon sections. Insets: zoom-in of a single colon crypt. Dotted lines: boundaries between MROIs: crypt apex (APEX), crypt mid (MID), sub-crypt (SUB-CRYPT), Peyer’s patch (PP), and muscularis interna (MI) (*Epcam, Tff3, Sma* scale bars: 200μm; *Cd48* scale bar 100μm). (**g**) Probability density (y axis) for posteriors over regional rate terms (*β*) inferred by the cSplotch model for the genes in (**f**) in each MROI (color code). (**h-j**) Validation of the cellular expression profiles predicted by cSplotch. (**h**) Pearson correlation coefficient (color bar) between mean snRNA-seq profiles (columns) and mean cSplotch *β′*s (rows) over the union of the top 50 cell type marker genes by snRNA-seq (**Methods**) for each of the top 8 cell types by mean cell fraction (columns, rows) in the sub-crypt for adult mice. Right: Pearson correlation for matching cell types. (**i**) Posteriors over cellular rate terms (*β*) inferred by cSplotch in CM in each abundant cell type (color code) for a goblet cell (*Prdx6*, top) and fibroblast (*Pdgfra*, bottom) marker in the CM region. (**j**) IF images of proximal colon sections for the genes (Prdx6 and *Pdgfra*) in (i). Insets as in (**f**). Scale bar: 200*μ*m.

To infer gene expression rates, cSplotch next uses these inferred cell compositions, MROI annotations, and sample covariates to fit a generalized linear model (GLM) of spatial gene expression across the atlas (**Extended Data Fig. 4, Methods**). The model performs Bayesian inference to apportion the aggregate expression of each gene in each spot to the contributing cell type(s) in the spot (characteristic expression rate (*β*); hierarchically formulated to account for covariate-driven variation), along with the effects of neighboring spots (through spatial autocorrelation (*ψ*)), and spot-specific random effects (*ϵ*) (**Methods**). Additionally, to integrate the spatial samples in the context of a common coordinate framework, we used the 14 MROI categories that were manually assigned by the histology at each spatial spot (**Methods**).

We validated cSplotch’s inferred expression rates in our colon aging atlas data. First, for highly expressed genes (∼10% of the genes in the study; detected in >45% of all spots; **Fig. 2b, red;** *e*.*g. Tff3, Ceacam1, Acta2*), the expression estimates of cSplotch were highly correlated to those obtained by transcripts-per-million (TPM) normalization, and the spatial autocorrelation and spot-level variation components of cSplotch captured a substantial portion of the variance (**Fig. 2c-e**). Second, cSplotch recovered the correct spatial differential expression patterns in each MROI compared to immunofluorescence (IF) staining for four selected marker genes: the highly expressed *Epcam* and *Tff3* (enriched in crypt apex (APEX)), *Acta2* (SMA; enriched in *muscularis interna* (MI)), as well as *Cd48*, which is expressed in only 2.7% of spots (enriched in the PP) (**Fig. 2f,g**). Third, the characteristic expression levels inferred for each cell type in a given MROI were most correlated with snRNA-Seq profiles from the expected cell type, for cell types present at both high (*e*.*g*. goblet cells) and low (*e*.*g*. fibroblasts) frequencies (**Fig. 2h**). Fourth, cSplotch correctly assigned *Prdx6* expression to goblet cells and *Pdgfra* to fibroblasts in the cross-mucosa (CM) region, confirming the accuracy of cSplotch in describing gene expression in a complex tissue region comprised of multiple cell types of varying proportions (**Fig. 2i,j**). Fifth, cSplotch’s accurately deconvoluted simulated ST data, generated by constructing spots using mixtures of snRNA-Seq profiles (**Extended Data Table 5, Methods**). cSplotch successfully reconstructed mean profiles for snRNA-seq (**Extended Data Fig. 5a**), morphological cell type clusters (**Extended Data Fig. 5b**), and differential expression patterns detected from snRNA-seq (**Extended Data Fig. 5c;** 94-100% agreement on DE genes), even when the cellular compositions used to deconvolve the simulated data were corrupted by Gaussian noise to reflect imperfect cell type annotations (**Extended Data Fig. 5a,b**). Finally, we estimated the impact of the number of individuals and tissues profiled on statistical power, by incrementally downsampling combinations of individuals (n=1,…,6 of 6) and tissue sections (n=2,…,8 of 52). Even when using one animal and two tissue sections, cSplotch captured meaningful expression signals, as reflected by the low KL divergence from the posterior distributions derived using full data for that animal’s covariate group (age and region), and accuracy improved significantly with at least four mice (**Extended Data Fig. 6a; Methods**). Increasing the number of mice improved the estimation independent of expression level, whereas increasing the number of tissue sections per mouse improved the estimation of lowly expressed genes (**Extended Data Fig. 6b; Methods**). In our full dataset, we sample at least five mice per time point, and a minimum of 9 tissue sections per mouse, exceeding these thresholds.

### Cell composition and cell type-specific expression across the proximal to distal colonic axis correlates with functional variation

To better describe the variation in tissue structure and function across the colonic tract, we first applied cSplotch to identify tissue-scale changes in cellular composition and spatial cellular gene expression along the proximal-distal axis in the adult (12 weeks) mouse colon. We analyzed 134 sections (43 proximal, 43 middle, 48 distal, 6,399 spots, ∼55 nuclei per spot) across 6 age-matched and gender-balanced mice (3 males and 3 females) (**Fig. 3a**). Relying on the classification in 14 MROIs, we estimated cell abundance and cell type-specific expression of 17 distinct cell types.

**Fig. 3.**
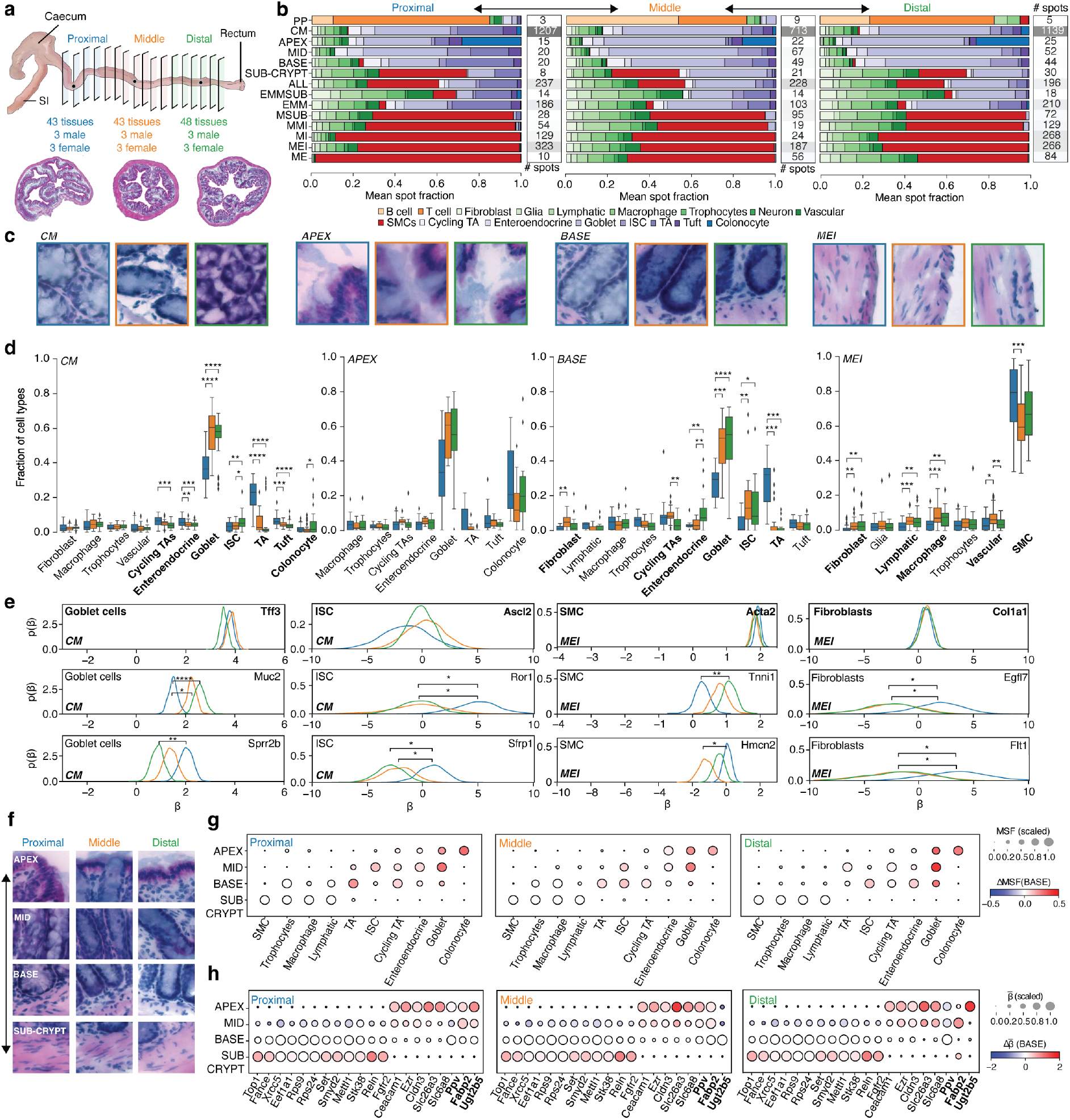
Regional differences in cell composition and function in the adult colon. **(a**) The atlas spans structural variation in the mouse colon along the proximal-distal axis. Illustrative H&E images of sections from proximal (left, blue), middle (mid, orange) and distal (right, green) regions. (**b**) Variation in cellular composition of MROIs across the proximal distal axis. Proportion of cells of each type (x axis, stacked bars; color code) in each MROI (y axis) in the proximal (left), middle (mid) and distal (right) regions of adult (12w) colon. # spots (right): Number of spots per MROI (color scale) in each colon region. (**c, d**) Variation in cell type composition of the same MROI type along the proximal-distal axis. (**c**) Representative H&E image patches from selected ST spots in four MROIs (left to right) from proximal (left, blue), middle (mid, orange) and distal (right, green) regions. (**d**) Fraction of cells (y axis) of each cell type (x axis) in the proximal (blue), middle (orange), and distal (green) regions in each of the MROI type in (c). Center black line, median; color-coded box, interquartile range; error bars, 1.5x interquartile range; *: 0.01 < FDR <= 0.05; **: 10^−3^ < FDR <= 0.01; ***: 10^−4^ < FDR <= 10^−3^; ****: FDR <= 10^−4^ (Welch’s *t*-test). Only cell types observed at a rate of 2% or greater across all spots in an MROI are shown. (**e**) Cell specific expression patterns vary across colon regions. Estimated posterior distributions of expression rate (*β*) for different genes in each colonic region (color) in goblet cells and ISCs in CM (two left columns) or SMCs and fibroblasts in MEI (two right columns). Bold: Canonical markers. Brackets: significant differential expression (*: Bayes factor (BF)>2; **: BF>10; ***: BF>30; ****: BF>100). (**f-h**) Variation in structure, cell composition and gene expression along locations. (**f**) Selected H&E image patches from ST across four MROIs (rows) from three colon regions (columns). (**g**) Scaled (dot size) and absolute (dot color) change in mean spot fraction (MSF, dot color) of each cell type (columns) in each MROI (rows) relate to BASE from three colon regions (panels). (**h**) Scaled (dot size) and absolute (dot color) change in mean log expression 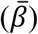 of each gene (columns) in each MROI (rows) relative to BASE from three colon regions (panels). Bolded genes: expression gradients.

Comparison of cell abundance and cell type-specific expression within individual MROIs between the proximal, middle and distal colon (**Fig. 3b**) showed that most cell type frequencies in most MROIs vary extensively along the regions, especially in the transition from the proximal to the middle colon (**Extended Data Table 6**). For example the abundance of intestinal stem cells (ISC), transit-amplifying (TA) cells, and goblet cells in the mucosa, and of smooth muscle cells (SMCs) and fibroblasts in the muscular layers changed significantly between the segments (BH-corrected Welch’s *t*-test, p<0.05) in each of four MROIs (CM, n = 3,059 spots; MEI, n = 776 spots; BASE, n = 113 spots; APEX, n = 62) (**Fig. 3c,d, Extended Data Table 6**). In all three crypt MROIs (CM, crypt base (BASE) and crypt apex (APEX)) goblet cell proportions increased and TA frequencies decreased from the proximal to the distal colonic segment, and in the non-apical crypt MROIs (CM and BASE) ISC proportions increased distally. In MEI (the layer spanning the *muscularis interna* and *externa*), SMC abundance declines distally, while fibroblasts, lymphatic, macrophage and vascular cells grow in frequency.

We also identified cell-type specific gene expression patterns that are associated (in conjunction with cell abundance) with the proximal to distal axis. cSplotch, analysis showed that canonical cell type-specific markers are similarly expressed across the three colonic regions, based on the posterior distributions of cellular expression rates (**Fig. 3e top, Extended Data Fig. 7, bold**), with stronger cell type-specificity signals than snRNA-seq alone (**Extended Data Fig. 8**) (*Tff3, Clca1* and *Lypd8* for goblet cells^56,57^; *Ascl2, Lgr5*, and *Ephb2* for intestinal stem cells^58–60^; *Acta2, Cnn1* and *Myh11* for SMCs^61^; *Col1a1, Thy1*, and *Postn* for fibroblasts^62^). In contrast, other genes had significant cell type-specific regional differences (Bayes factor > 2) across the proximal-to-distal axis (**Fig. 3e**, middle and bottom, **Extended Data Fig. 7** non-bold, **Methods**). For example, in the CM, goblet cells upregulated the antimicrobial genes *Sprr2b* and *Sprr2a3*^*63*^ proximally, and the key gel-forming mucin *Muc2* distally. Such goblet-specific expression patterns are consistent with the role of the proximal colon in controlling microbial fermentation, and the requirement for a thick mucus layer as a protective barrier distally^64^. Within the same MROI, ISCs expressed proximally higher levels of canonical Wnt inhibitors (*e*.*g*., *Sfrp1*) and mediators of the non-canonical pathway (*e*.*g*., *Ror1, Wnt4*). As the strength of canonical Wnt signaling is tightly regulated locally and associated with ISC self-renewal^65^, the proximal colon may restrain this pathway and experience a slower cellular turnover. Conversely, the distal colon may require less restrained canonical Wnt signaling to replenish an expanded goblet population, which has the shortest life span across mucosal cell types^66^. In MEI, SMCs upregulated *Hmcn2* proximally and *Tnn1* distally. As *Hmcn2* is associated with Hirschsprung’s disease^67^, and *Tnn1* helps enforce contractility^68^, their regional-specific expression can support autonomous peristalsis proximally, and voluntary contraction distally. In MEI, genes encoding for repressors of vasculogenesis and lymphogenesis, like *Egfl7, Flt1*, and *Mdfic*^*69,70*^, were upregulated by the fibroblasts proximally, indicating the loss of such mediators as potential mechanism for the expansion of the vascular system distally, where it is required for water reabsorption.

Collectively, we found that the variation in both cell abundance and cell-type specific gene expression along the colon longitudinal axis correlates with distinct digestive functions along the colonic tract.

### Cell type and gene expression gradients along the crypt axis associated with colonic regeneration and functional differentiation

The intestinal crypt is responsible for the continuous regeneration of the highly specialized colonic mucosa^3^. As ISCs divide at the bottom of the crypt, they migrate towards the apex differentiating into the main specialized lineages (goblet and colonocytes), as well as a variety of rarer cell types, including enteroendocrine and chemosensory cells. A morphogen gradient modulates the balance between epithelial proliferation and differentiation, but its composition at different biological scales is not fully characterized^71^. To date, targeted imaging approaches have characterized only a limited number of positionally-restricted signaling mediators regulating crypt structure and function^72^.

Both cell morphologies and identities displayed distinct zonation patterns across four MROIs of the crypt axis (sub-crypt (SUB-CRYPT), BASE, crypt mid (MID), and APEX) (**Fig. 3f,g**). ISCs, TA, and cycling TA cell proportions decrease gradually from the crypt base to apex, goblet cell proportions follow an opposite gradient, colonocyte abundance peaks sharply in the apex, and enteroendocrine cells are evenly scattered across the crypt axis. SMCs are almost entirely confined to the subcrypt, while lymphatic cells, trophocytes, and macrophages gradually decline in a base-to-apex direction. These cell-type specific distributions are consistent with distinct positional dependencies. Colonocytes and SMCs are localized in close proximity to physical barrier domains (*e*.*g*., the colonic lumen and the lamina propria), while the distribution of epithelial cells (ISCs, TAs, goblets), immune cells (macrophages) and some mesenchymal cells (trophocytes, lymphatics) is consistent with a reliance on overlapping signaling cues from the opposite crypt poles. Finally, the homogeneous scattering of enteroendocrine cells may reflect the distribution of enteric nerve fibers.

At the molecular level, 321 genes had significant, monotonic variation across the crypt regardless of the colonic region (**Fig. 3h, Extended Data Table 7, Methods**). Some of these genes were significantly associated (BF > 2, l2fc > 0.5) with specific cell types in a given spatial niche. For example, many of the genes upregulated in BASE were both expressed in ISCs and implicated in key biosynthetic pathways, including DNA replication and repair (*Top1, Fance, Xrcc5*), protein synthesis (*Eef1a1, Rps9, Rps24*), and transcriptional (*Set, Smyd2*), translational (*Mettl1*) and posttranslational (*Stk38*) regulation (**Extended Data Table 8**). Genes upregulated at the apex were associated with the establishment of an apical polarity domain at the plasma membrane of goblet cells, including cytoskeletal (*Ezr*), cell-cell adhesion (*Ceacam1, Cldn3*), and transmembrane transport (*Slc26a3, Slc6a8*) genes (**Extended Data Table 8**). Additionally, receptors (*Fgfr2*) and ligands (*Reln*), mediating the cross-talk between ISCs^73–75^ and the underlying stromal compartment (*e*.*g*., lymphatics and trophocytes), were upregulated in the SUB-CRYPT and BASE. Another 118 genes had crypt-oriented gradient expression preferentially in one colonic region (23 proximal, 53 middle, 42 distal). These included genes related to key metabolic functions, including the detoxifying enzyme *Ugt2b5*, which is downregulated in experimental models of colitis^76^, the fatty-acid binding and importer protein *Fabp2*^*77*^, and the gut hormone *Ppy*, known to regulate food intake^78^ (**Fig. 3h**). These further show how each of three colonic regions may rely on partially distinct molecular gradients associated with different digestive functions.

In summary, our analysis revealed cell type-specific distributions, likely reflecting distinct positional dependencies, and identified a subset of pan-colon or region-specific gradient genes to prioritize mechanistic experiments on the crypt’s structure and function.

### Spatiotemporal variability of cell type abundances during colon aging

In the elderly human population, the large intestine is affected by common pathological conditions including constipation^79^, diverticulitis^80^, malnutrition^81^, a markedly increased risk of colorectal cancer^82^, and other features of intestinal senescence^8,18,83–89^. Yet, aging is a loosely defined condition, which is asynchronously experienced across populations with vastly variable phenotypic impact. Previous studies carried out on small cohorts, few time points, and using destructive methodologies have led to potentially conflicting findings. For example, both loss and gain of secretory cells have been reported to occur in the aging mouse intestine^18,88,90^.

To characterize intestinal cells during mouse aging, we tracked changes in the proportion of the most abundant cell types (ISCs, TA cells, goblet cells, and colonocyte) in the four crypt MROIs in each of the colonic regions over time (**Extended Data Fig. 9, Extended Data Table 9**). During the juvenile (up to 4 weeks) and adult (4 to 12 weeks) phases, there was limited variation in the ISC fraction over time in the SUB-CRYPT, BASE and MID, consistent with the highly regenerative nature of the colon, during both development and at steady-state. In the same compartments, TAs displayed a transient increase during the first four weeks after birth, followed by a decline between 6 and 12 weeks, as they increasingly transitioned towards the secretory lineage. Throughout juvenile and adult life, goblet cells steadily increased across all MROIs, reaching their peak number between 8 and 12 weeks. Although the structure and function of colonocytes is impacted by weaning, involving a nutritional transition from lactation to solid food starting 3 weeks after birth^91–93^, their relative proportion progressively decreased in the APEX up to 12 weeks, concomitantly with the increase in goblet cell fraction.

Importantly, there were multiple significant changes in cell composition of the same region between full reproductive maturity (12w) to end of life (2yr) across the crypt axis in the distal and middle colon, as well as, to a lesser extent, in the proximal region (**Extended Data Fig. 9**, shaded areas, **Extended Data Table 9**). Goblet cell frequency declined from young adult (12w) to aged (2yr) animals in all MROIs defining the crypt axis in the mid colon; in the SUB-CRYPT, BASE and MID regions in the distal colon, and in the BASE of the proximal colon. Conversely, the frequency of progenitor cells (ISCs and TAs) increased in old mice (2yr) in both the middle and distal colon, with TAs increasing with age across the entire crypt axis in the middle colon, and ISCs increasing with age in the SUB-CRYPT, BASE and MID MROIs in the distal colon. In the APEX of these segments (middle and distal), there was also a marked increase in colonocytes during that period. Hence, cell frequencies are substantially remodeled as animals age, with colonocytes progressively displacing a dwindling goblet population in the APEX, which may be especially impactful in the mid and distal mucosa, which rely more heavily on secretory cells at steady state. Goblet cells decline despite the concurrent increase in the proportion of ISCs and TAs in the SUBCRYPT, BASE, and MID. These temporal dynamics suggest an imbalance between progenitor proliferation and secretory differentiation, providing a cellular base for the defective regenerative function observed in the elderly.

### Variation in cell type abundances associated with activity of multicellular programs during colon aging

Next, to gain insights into the sequence of molecular events driving the change of mucosal cell identities in the senescent colon, we applied a unified quantitative framework for the identification of cellular and expression alterations. We used DIALOGUE^54^ to infer MCPs of genes with correlated expression patterns across multiple cell types over time (**Fig. 4a,b, Methods**). Specifically, each MCP consists of sub-sets of genes whose mean expression in one cell type is positively or negatively correlated with that of (potentially other) genes in one or more other cell types across time. Several of the MCPs with the highest degree of correlation between member genes had near-monotonic changes during the aging window (*e*.*g*., between 6 months and 2 years of age) in at least one colonic region (**Fig. 4c, Extended Data Table 10**).

**Fig. 4.**
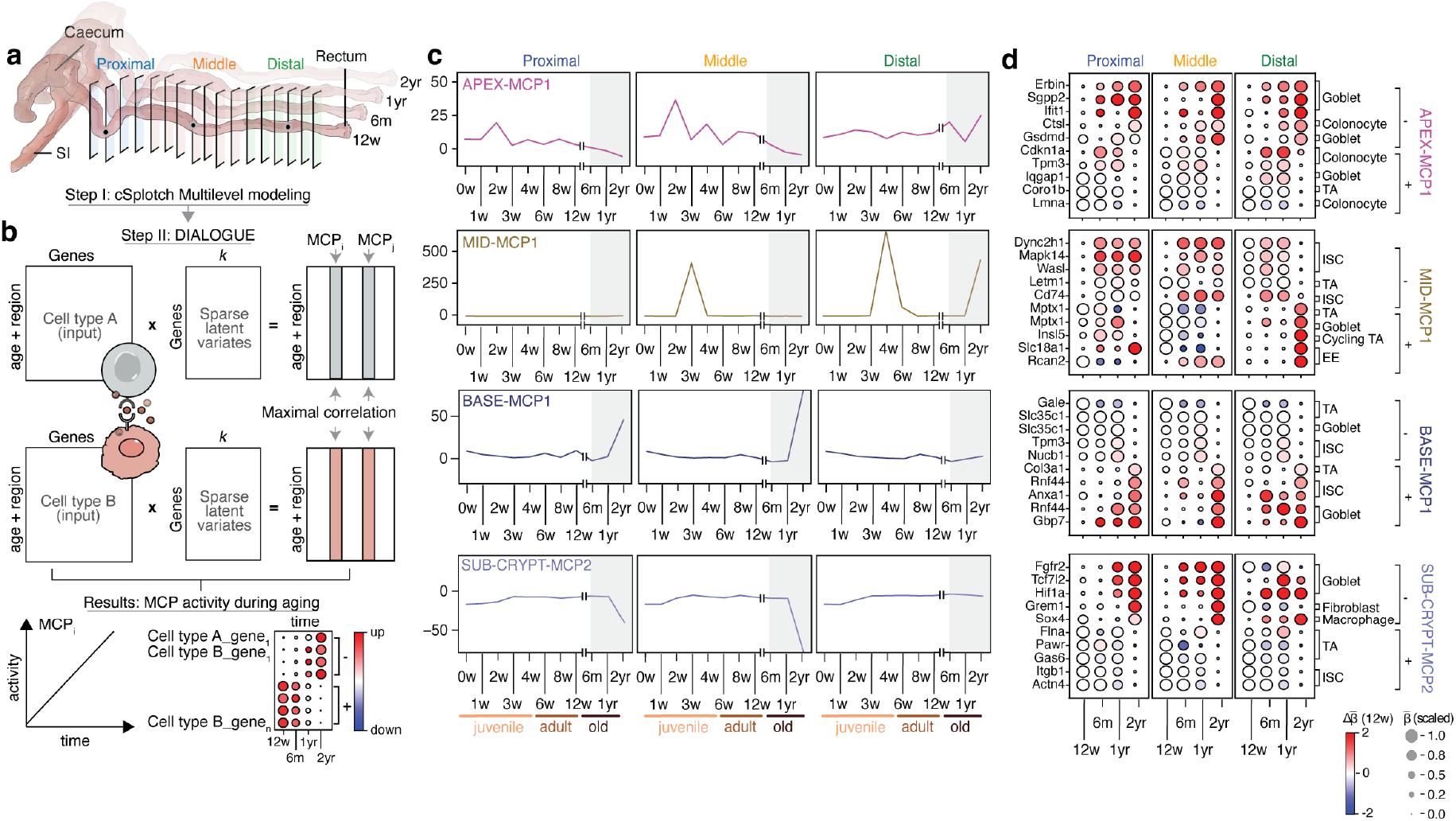
Spatiotemporal changes in cell composition and function during colon aging. **(a,b**) Analysis approach. cSplotch is used to characterize variability in cell-type specific gene expression across age and region (a), followed by MCP analysis with DIALOGUE^54^ (**b**) on inferred profiles for abundant cell types (e.g., cell types A and B) in each MROI across ages and regions. MCPs are returned as sets of genes with high correlation between two or more cell types across conditions (with positively (+, up) and negatively (-, down) contributing genes from each cell type (**Methods**). (**c,d**) Aging-associated MCPs. (**c**) Activity score (y axis) of selected MCPs (panels) from different MROIs (top left label) in each time point (x axis) and region (labels on top). Gray area: aging window. (**d**) Scaled (dot size) and absolute (dot color) change in log expression 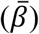 relative to 12 weeks of each gene (rows) in each time point (columns) for selected up (+) and down (-) regulated genes from the MCPs in (**c**), sorted by their associated cell type (label on right).

SUB-CRYPT-MCP2, whose activity markedly declines at one year of age in the proximal and middle colon (**Fig. 4d**), may help explain some of the substantial changes in cell composition with age in the SUB-CRYPT region, where goblet cells decline and progenitors increase at 2 years of age (**Extended Data Fig. 9**). Specifically, SUB-CRYPT-MCP2, consists of genes from nine cell types, epithelial (ISCs, TAs, cycling TAs, goblets), macrophages, trophocytes, lymphatic, vascular, and SMCs. These included age-increasing pro-inflammatory genes in macrophages (*Sox4*)^94^, and signaling mediators in goblet cells (*Fgfr2, Tcf7l2, Hif1a*)^95–97^ and trophocytes (*Grem1*)^98^; many of which are involved in epithelial dedifferentiation^95,96,99^, inflammation^97^, and malignant transformation^98,100,101^, especially colitis-associated^100,101^ (**Extended Data Table 11**). Age-declining genes included intestinal mechanotransduction (*Itgb1, Flna, Actn4*)^102–104^, cell death regulation (*Gas6, Pawr*)^105,106^, and tumor suppressor genes (*Itgb1, Flna, Gas6, Pawr*)^102,105,107,108^ (TSG) expressed in ISCs and TAs (**Fig. 4d**). As the adult distal SUB-CRYPT had enhanced WNT signaling (**Fig. 3e**), the loss of such TSGs could explain a shift in the balance between self-renewal and differentiation, leading to an expanded stem cell compartment and defective lineage priming. In contrast, the adult proximal SUB-CRYPT was enriched for canonical Wnt inhibitors. Here, TSG loss alone may be insufficient to drive a wider progenitor compartment in the absence of a permissive microenvironment. Instead, we observed an upregulation of regenerative signaling mediators (*Fgfr2, Tcf7l2, Hif1a*), which may eventually promote dedifferentiation of the secretory lineage into progenitor cells. In line with such a hypothesis, the middle colon displayed an intermediate scenario where the rewiring of goblet signaling was associated with an expansion of the TA fraction.

The activity of the BASE-MCP1 program, which consists of gene expression in ISCs, TAs and goblet cells, increases in the proximal and middle colon following one year of age. At two years of age, this MROI displayed loss of goblets throughout the colon, and a gain in progenitors in the middle and distal colon. The BASE-MCP1 genes increasing with age include the key fetal stem cell marker *Anxa1*^*109*^, upregulated in ISCs and associated with damage-induced regeneration^110^; *Rnf44*^*111*^ and *Gbp7*^*112*^, known to be induced by inflammation and expressed in ISCs and goblets, and the pro-tumorigenic extracellular matrix protein *Col3a1*^*113,114*^ found in TAs (**Fig. 4d**). Age-declining genes include metabolic regulators of glycosylated surface proteins and lipids, like *Slc35a1*^*115*^ and *Gale*^*116*^, important for immune recognition and pathogen infection, expressed in TAs and goblets, and *Tpm3*, a gene recurrently lost in CRC^117^, expressed in ISCs. The progressive activation with aging of damage-induced regeneration genes (*Anxa1, Rnf44, Gbp7*) in a distal-to-proximal colon gradient, may suggest that damage-induced responses become pervasive with aging, predisposing the colon to transformation.

The MID-MCP1 program, with genes expressed in ISCs, TAs, goblets, enteroendocrine and tuft cells, had different aging-related activity in the proximal and middle *vs*. distal colon. At two years, goblet cells are lost and progenitors increase in the middle and distal MID MROI, while no significant cell composition changes are observed in the proximal region. MID-MCP1 genes with neuroendocrine and metabolic functions (*Insl5*^*118*^, *Slc18a1*^*119,120*^, and *Rcan2*^*121,122*^) declined with aging in the proximal and middle colon, but increased in the distal colon. This suggested aging-related anteriorization of the longitudinal colonic patterning, an event also associated with the emergence of malignant states^123,124^. Such reprogramming was additionally coupled with the change in expression of genes involved in mucosal inflammation (*Cd74, Dync2h1*)^125,126^, stress response (*Mapk14, Letm1*)^127,128^ and CRC initiation (*Wasl*)^129^.

Finally, the activity of the APEX-MCP1 program, which includes genes expressed in TAs, colonocytes and goblets, declined across the colonic tract during aging (**Fig. 4c**). MCP genes upregulated with age included inflammation genes related to pyroptosis (*Gsdmd*)^130^, proteolysis (*Ctsl*)^131^, membrane integrity (*Sgpp2*)^132^, cell polarity (*Erbin*)^133^, and the DNA damage response (*Ifit1*)^134^. APEX-MCP1 genes downregulated with aging included cytoskeleton regulators (*Coro1b, Iqgap1, Tpm3*)^117,135,136^ expressed in TAs, colonocytes, and goblet cells, and *Lmna*, an aging marker expressed in colonocytes encoding the nuclear lamin A/C and mutated in congenital premature aging syndromes^137,138^, and *Cdkn1a* (p21)^139^ (**Fig. 4d**). Thus, the aging colonic APEX experienced a diffuse inflammatory state, associated with the expression of canonical aging markers and the downregulation of cell cycle regulators, indicating the acquisition of a senescent state in the apical colonocytes coupled with the loss of tumor suppressor mechanisms.

## DISCUSSION

Characterizing spatial patterning in large organs requires us to harmonize and relate molecular and morphological profiles measured by single cell and spatial RNA-seq methods. Despite significant progress to address spatial variance in expression and histological features, existing analysis methodologies are still mostly applied to a single tissue section at a time. As a result, they may fail to identify generalizable tissue function or identify how key features vary along anatomical, functional or temporal axes. Here, we developed a comprehensive experimental and computational framework and used it to present the first systematic atlas of colon aging.

Our analysis revealed changes in cell composition and cell-type specific gene expression across multiple inter-leaving scales: gross anatomical scale (the longitudinal proximal-distal axis), fine histological scale (the crypt base to apex axis), and the temporal scale (ages from newborn to old). Along the proximal-distal axis these variations correlate with distinct digestive functions along the colonic tract. Along the crypt axis, we revealed cell-type specific positional patterns, and genes with either pan-colonic or region-specific spatially-restricted expression across the main crypt axis. The positionally-oriented balance between progenitor self-renewal and lineage differentiation was associated with a base to apex gradient between biosynthetic and cytoarchitectural regulators, prioritizing pathways and genes for functional validation. Along the temporal axis, aging was associated with a pervasive loss of goblet cells and the establishment of a diffuse inflammatory state, spanning expression programs in multiple cells, across the colon, and revealed the specific impact of such a scenario at different levels of the crypt axis and in distinct regions of the colonic tract. In the upper parts of the crypts, the progressive displacement of goblet cells by apical colonocytes, or the anteriorization of the colonic metabolic patterning, can provide molecular insights towards a mechanistic understanding of common geriatric conditions, like constipation and malnutrition, paving the way for interventions aimed to support the quality of life, and preventing systemic consequences. At the bottom of the crypt, increasing levels of damage coupled with persistent inflammation are associated with loss of tumor suppressor gene expression, excessive expansion of the progenitor compartment, and cellular dedifferentiation towards fetal-like states. Such temporal dynamics are consistent with the elevated incidence of CRC premalignant states in the elderly^140^, and illuminate specific cell types and genes for guiding preventive and diagnostic strategies.

Using the canonical system of the mouse colon, we have demonstrated how contextual spatial and temporal information can help decipher large-scale molecular datasets, and how statistical models like cSplotch can be used to connect tissue architecture with the pathological alterations leading to aging and disease. We believe that this model framework has utility in a wide range of tissue systems, and hope that it may help to bridge the gap between single-cell and spatial transcriptomic studies.

## METHODS

### Murine tissue collections

C57BL/6J mice were obtained from The Jackson Laboratory (Bar Harbor, ME) and maintained in accordance with ethical guidelines monitored by the Institutional Animal Care and Use Committees (IACUC) established by the Division of Comparative Medicine at the Broad Institute of MIT and Harvard and Columbia University, and consistent with the Guide for Care and Use of Laboratory Animals, National Research Council, 1996 (institutional animal welfare assurance no. A4711-01), with protocols 0122-10-16 and AABI3617, respectively. Colons were collected within 5 min of death and flushed using ice-cold 1x PBS (Gibco) using a straight gauge needle with a 2.4mm tip (Kent Scientific Corporation, USA) for mice >3 weeks of age or a newborn feeding needle (Cadence Science, USA) for mice ages 3 weeks. The rest of the mice were processed without colon flushing prior to freezing. Tissues were then dried and embedded in Optimal Cutting Temperature (OCT, Fisher Healthcare, USA) in large molds (VWR, USA). Samples were then plunged onto a metal plate pre-chilled and sitting on top of dry ice for 2min or until complete freezing. Samples were transferred and stored at −80°C until use. Cryosections were cut at 10μm thickness onto ST slides, and stored at −80°C for at most 2 days. For nucleus extractions, colons were separated from the animals within 5 minutes of death and each colon was separately flushed similar to collecting tissue for ST. Approximately 0.5cm of tissue from three different colon regions was placed on a tissue and dried before transfer to a sterile dish to ensure all excess water was gently removed from the sample. Tissues were then placed in a 1.5 mL tube on dry ice and, upon freezing, transferred to −80°C until subsequent tissue processing.

### Immunostaining and epifluorescent microscopy

A superfrost slide (ThermoFisher Scientific, USA) with a mounted tissue section was dried to 37°C for 4 minutes on a thermal incubator (Eppendorf Thermomixer Option C, Germany) followed by *in situ* fixation in 4% PFA (Sigma Aldrich, USA) at RT and washing in 50mL 1x PBS (Gibco). Slides were placed in a chamber holder (ProPlate Multi-Array slide system; GraceBioLabs, USA) to allow for incubations in predefined conditions and volumes. All following antibody incubations were performed at 4°C. To block tissues from nonspecific antibody binding, 1:100 TruStain FcX™ PLUS (anti-mouse CD16/32, Biolegend, USA) antibody in 1x Perm/Wash buffer (BD, USA) was added and tissues were incubated for 30min. Tissues were washed 3 times with 1x PBS-T (0.05% Tween-20, Sigma, USA). Antibodies were added at a 1:100 dilution in 1x Perm/Wash buffer and incubated for 30min. If an unlabeled primary antibody was used, tissues were again washed and stained with a labeled secondary antibody prepared in 1x Perm/Wash buffer (BD, USA). Tissues were then washed in the same fashion before counterstaining with Hoechst (10mg/ml, ThermoFisher Scientific, USA) diluted 1:1000 in 1xPBS (ThermoFisher Scientific, USA) for 5 min. This was followed by another wash cycle after which slides were air dried and mounted with 85% glycerol prior to imaging. Primary antibodies were diluted and clone and provider information were as follows: EPCAM (Biolegend, Alexa-647 labeled primary, clone G8.8, 1:100 dilution), TFF3 (Abcam, unlabeled primary, EPR26048-14, 1:100 dilution), SMA (Abcam, unlabeled primary, clone EPR5368, 1:100 dilution), FABP4 (Abcam, Alexa-647 labeled, clone EPR3579, 1:250 dilution), FABP7 (Abcam, unlabeled primary, clone EPR24033-13, 1:100 dilution), CD48 (Biolegend, APC labeled, clone HM48-1, 1:100 dilution), PRDX6 (Abcam unlabeled primary antibody, clone EPR3754, 1:500 dilution), CD74 (Biolegend unlabeled primary antibody, clone In1/CD74, 1:250 dilution), PDGRFA (Abcam unlabeled primary antibody, clone EPR22059-270, 1:1000 dilution), goat anti-rat IgG (Life Technologies, Alexa Fluor 488 Goat anti-rat IgG, 1:100 dilution), donkey anti-rabbit IgG (Biolegend, Alexa 647 labeled, clone Poly6064, 1:100 dilution). Epifluorescent images were acquired on an Axio Imager Z2 microscope using a PhotoFLuor LM-75 lightsource (89North, USA) in combination with a Plan-APOCHROMAT 40x/0.75 objective (Carl Zeiss, Germany). Images were stitched using Vslide (v1.0.0, MetaSystems GmbH).

### Slide production

Spatially barcoded arrays were produced as previously described^31,141^, with six active surfaces per slide. Using Codelink chemistry (Surmodics, USA), 5’ amine-modified DNA oligonucleotides (5’-[AmC6]dUdUdUdUd-[Illumina_adaptor]-[spatial barcode]-[UMI]-[20T]-VN) (IDT, USA) were bound to the slide surface using 100pL droplet deposition (ArrayJet LTD, Scotland, UK), This patterning formed 100μm spots with a 200μm spot-to-spot pitch distances, for 1,007 spatially barcoded spot conjugations in a 6.2mm x 6.6mm capture area. Finally, the unconjugated surface was blocked using a blocking buffer at 50°C (50 mM ethanolamine, 0.1 M Tris, pH 9) for 30min before washing the slides in 4x SSC, 0.1% SDS (pre-warmed to 50°C) for 30 min, rinsing them with deionized water and drying.

### Histology for Spatial Transcriptomics

Tissue sections were adhered to the ST arrays at 37°C for 1 min, *in situ* fixated in 4% PFA (Sigma Aldrich, USA) at RT and washed in 50mL 1x PBS (Gibco). In most cases, 4 tissue sections were fitted onto one ST active area. Tissues were then dried for 1min in isopropanol, followed by hematoxylin and eosin (H&E) staining. Briefly, tissue sections were exposed to 100% Mayer’s hematoxylin (DAKO, Agilent) for 6 minutes followed by washing for 2 min in deionized water at RT. To adjust the pH, slides were briefly dipped in DAKO’s Bluing buffer (Agilent) and then counterstained in 5% eosin diluted in Tris-AA (pH 7.2) for 1 min. Slides were again washed in deionized water and dried prior to mounting them with 85% glycerol prior to imaging. All samples were imaged on an Axio Imager Z2 microscope equipped with a 20x/0.8 Plan-APOCHROMAT (Carl Zeiss Microscopy, Germany) and the resulting images stitched with Vslide (v1.0.0, MetaSystems GmbH).

### In situ and library preparation of Spatial Transcriptomics reactions

After imaging, coverslips were removed in deionized water and permeabilization reactions immediately started. First, tissue samples were permeabilized with 120μl reagent per reaction of collagenase I (200U) in 1x HBSS (both from ThermoFisher Scientific, USA). Pre-permeabilization reagent removal was followed by a 180μl wash in 0.1X Saline Sodium Citrate (SSC, Sigma-Aldrich, USA) at 37°C. Next, tissue was permeabilized with 75μl 0.1% pepsin (pH 1, Sigma-Aldrich, USA) at 37°C for 10min, followed by another wash with 0.1x SCC. Reverse transcription (RT) was performed by the addition of 75μl RT reagents: 50ng/μl actinomycin D (Sigma-Aldrich, USA), 0.5mM dNTPs (ThermoFisher Scientific, USA), 0.20μg/μl BSA, 1X First strand buffer, 5mM DTT, 2U/μl RNaseOUT, 20U/μl Superscript III (all from ThermoFisher Scientific, USA). Tissues were digested from the slide surface by 1h incubation in proteinase K (Qiagen, Germany) at 56°C. Slides were washed as suggested by the slide manufactures (Codelink, Surmodics): 10min 2x SCC with 0.1% SDS (Sigma Aldrich), 1min 0.2x SCC and 1min 0.1X SSC. To remove spatial cDNA:mRNA hybrids from the array surface, Uracil-Specific Excision Reagent (NEB, USA) was used as previously described^142^.The reaction was run for 2h at 37°C and the resulting spatially barcoded cDNA libraries were collected and libraries prepared as described previously^142^. Briefly, cDNA:RNA hybrids collected from the array surface were used as input in the first part of library preparation reactions. RNA strands were digested and used as primer to make dsDNA using DNA Polymerase I and RNaseH (2.7X First strand buffer, 3.7 U/μl DNA polymerase I and 0.2 U/μl Ribonuclease H (all from ThermoFisher Scientific, USA)) for 2h at 16°C. The material was made into blunt-end dsRNA products with 15U T4 DNA polymerase (NEB, USA) for 20 minutes at 16°C and reactions stopped by addition of 20mM EDTA (pH 8.0, ThermoFisher Scientific, USA). dsDNA was purified using Ampure XP (Beckman Coulter, USA) at a bead to cDNA ratio of 1:1. Next, the material was linearly amplified using a T7 promoter sequencing initially embedding in the oligonucleotides on the array surface by adding 27.8μl of the T7 reaction mix (46.2mM rNTPs, 1.5X T7 reaction buffer, 1.54 U/μl SUPERaseIN inhibitor and 2.3U/μl T7 enzyme; all from ThermoFisher Scientific, USA) for 14h at 37°C. This was followed by a bead cleanup with RNAclean Ampure XP beads (Beckman Coulter, USA) at a beads:aRNA ratio of 1.8:1. 8μl of the eluted aRNA was used as input to the following reactions. 2.5μl of 3μM aRNA adapters [rApp]AGATCGGAAGAGCACACGTCTGAACTCCAGTCAC[ddC] were added to 8μl of aRNA. The reaction was then incubated at 70°C in a PCR machine for 2min and immediately chilled on wet ice. To ligate the adaptors, 4.5μl T4 RNA ligation mix (3.3X T4 RNA ligase buffer, 66U/μl truncated T4 ligase 2 and 13U/μl murine RNAse inhibitor (all from NEB, USA)) were added at 25°C for 1h, followed by an Ampure XP (Beckam Coulter, USA) bead purification at a bead:cDNA ratio of 1.8:1. 1:1 v/v of 20μM GTGACTGGAGTTCAGACGTGTGCTCTTCCGA and 10mM dNTPs were added to the ligated samples and heated to 65°C for 5min. Reverse transcription took place by adding 2.5X First strand buffer, 13mM DTT, 5 U/μl RNaseOUT and 25 U/μl Superscript III (all from Thermo Fisher Scientific, USA). Samples were incubated at 50°C for 1h. 10μl of nuclease-free water were added followed by a final Ampure XP bead purification at bead:cDNA ratio of 1.7:1 with a final elution of 10μl nuclease-free water. qPCR library quantification and indexing were performed as previously described^142^.

### Nucleus extraction and library preparation for snRNA-seq

SnRNA-Seq was performed as previously described^17,143^. Specifically, frozen tissues were taken from 1.5ml tubes used for storage at −80°C and placed in pre-chilled 1 mL extraction buffer (0.03% Tween-20, 146mM NaCl, 10mM Tris pH 8.0, 1mMM CaCl2 and 21mM MgCl2; all from Sigma Aldrich, USA), supplemented with 400U RNasin Plus inhibitor (Promega Corporation, USA) and 200U SUPERaseIn RNase Inhibitor (ThermoFisher Scientific, USA). Tissues were disintegrated by chopping with tungsten Carbide Straight 11.5 cm Fine Scissors (14558-11, Fine Science Tools, Foster City, CA) for 10 minutes on ice. To avoid clogging, samples were filled through a 40 μm strainer (Falcon). To release any leftover material from the strainer, it was cleaned with the extraction buffer without the addition of Tween-20, followed by centrifugation to pellet the nuclei at 500g for 5 mins at 4°C. Supernatants were removed and nuclei were resuspended in a 100μL extraction buffer without the addition of Tween-20 before filtering through a 40 μm strainer-capped into a round bottom tube (Falcon). Nuclei were counted and ∼8,000 nuclei were loaded per channel on the GemCode Single Cell Platform using the GemCode Gel Bead kit, Chip and Library Kits (10X Genomics, Pleasanton, CA), following the manufacturer’s protocol. Briefly, nuclei were partitioned into Gel Beads in Emulsion (GEMs), lysed and barcoded using reverse transcription reactions, followed by amplification, shearing and 5′ adapter and library indexing.

### Spatial Transcriptomics sequencing and demultiplexing

ST cDNA libraries were diluted to 4nM and 1.08pm libraries loaded for sequencing on an Illumina NextSeq 550 (Illumina, USA) using paired-end sequencing (R1 30bp, R2 55bp). Samples were sequenced at a mean depth of 65.6 million paired-end reads depth. fastq reads were generated with bcl2fastq2. ST Pipeline^144^ was used to process the resulting fastq files. Briefly, 5nt trimmed R2 was used for mapping to the mouse genome using STAR^145^. After that, mapped reads were annotated using HTseq-count^146^. Spatial barcodes were collapsed using TagGD^144,147^ modified demultiplexer (k-mer 6, mismatches 2). Then, unique molecular identifiers (UMIs) mapped to the same transcript and spatial barcode were collapsed using naive clustering with one mismatch allowed in the mapping process as described in umi-tools^148^. The output genes-by-barcode matrix was used in all further processing steps. Average library saturation was 78.3%. To focus on reliably detected genes across spots, genes detected in fewer than 2% of spots were removed, as were spots with fewer than 100 UMIs across all genes. The resulting median number of genes and UMI transcripts per spatial spot was 2,164 (10^th^ percentile was 829 and 90^th^ percentile was 4,370) and 4,343 (10^th^ percentile was 1,246 and 90^th^ percentile was 13,055).

### snRNA-seq sequencing and demultiplexing

Libraries were sequenced on an Illumina NextSeq 550 (R1: 26 bases; R2: 55 bases) or a NovaSeq 6000 (R1: 28 bases; R2: 94 bases). CellRanger v3.0 was used for all initial data pre-processing. Fastq reads were first demultiplexed and then mapped to the reference mm10 transcriptome, augmented to allow for counting of all transcript tags in addition to counting exons as suggested by 10X Genomics. Each barcode was connected to a particular cell and UMIs were collapsed to account for duplicated transcripts. Filtered matrices reflecting digital gene expression (DGE) for each sample and cell were extracted from the pipeline.

### Analysis of snRNA-seq colon data

DGE matrices were concatenated from samples collected at 10 different ages. Potential doublets were removed using scrublet^149^ (∼12% of barcodes in the dataset). To facilitate downstream analyses, all snRNA-seq data were combined with the mouse colon droplet data from Drokhlyansky *et al*^*17*^ (also collected by our lab). Nucleus profiles with at least 800 genes expressed in a minimum of 10 cells were kept for further analysis. To ensure that only highest quality profiles were retained, profiles with less than 800 UMIs or more than 30% mitochondrial or ribosomal transcripts were also removed. Data were then normalized by the total number of transcripts or UMIs per nucleus profile and converted to transcripts-per-10,000 to account for differences in sequencing depth. Data were regressed out based on genes listed as differentially expressed in Drokhlyansky *et al*^*17*^ with the following cut offs: log_2_(fold change)>1 and Benjamini-Hochberg (BH) FDR<0.01 (Likelihood ratio test). Briefly, these genes were chosen based on the following criteria: mean (μ) and coefficient of variation (CV) of expression were calculated ofr each gene and partitioned into 20 equal-frequency bins. LOESS regression was used to fit the relationship between log(CV) and log(μ). Genes with the 1,500 highest residuals were equally sampled across these bins. To account for differences in batches, this was performed for each sample separately and a consensus list of 1,500 genes with the highest recovery rates was selected. Additionally, to account for cycling cells, both cell cycle scores (as in scanpy.tl.score_genes_cell_cycle^150^) and mitochondrial content scores in each cell were regressed. All of the following processing steps including clustering were performed with scanpy^150^. Overall, final analyzed data included 403,797 nucleus profiles with an average 2,281 genes and 4,305 UMIs per nucleus.

Batch correction was performed with Harmony^151^ as follows. Dimensionality reduction was performed using principal components analysis (PCA) and then a *k*-nearest neighbors (*k*-NN) was constructed using estimated k=20 neighbors and the first 40 PCs. Convergence was reached after 10 iterations and the Harmony corrected reduced d^150^ data was then clustered using Phenograph^152^ with *k*=25 nearest neighbors using the Minkowski metric. After clustering, cell type labels from Drokhlyansky *et al*^*17*^ were manually transferred to annotate clusters.

### Image and Spatial Transcriptomics data pre-processing

H&E images were processed with SpoTteR^153^. Briefly, original H&E images were scaled to approximately 500×500 pixels. Then, the tissue section was masked generously from the image through 10% quantile thresholding in a user-defined color channel. To detect probable spot centers, the image Hessian was computed. The spot centers then acted as potential grid points that were likely part of a regular grid structure and were selected by calculating the x and y distances between all detected centers. A regular grid was then fitted to these potential grid points using a custom optimizer based on the *nlminb* function of the R package stats, which minimizes the distance of potential grid points to the suggested regular grid while assuming angles of 90° and 42 starting grid points per row and column. Trough an iterative process, in which the 0.1% potential grid points that least fit the grid were removed in each iteration, the number of grid points per row and column were updated, and a new grid was fitted until the target number of grid points per row (here 35) and column (here 33) were reached. Finally, those grid points that overlapped the tissue sections were identified by building a mask that represented the tissue area and registering all grid points that were present in this mask. In case a sectioning artifact was present, the corresponding ST spot was removed from all subsequent analyses.

### Spatial Transcriptomics spot annotation

To assign each ST spot with a corresponding histological tag, a previously described cloud-based interface^33^ was used to assign each spot (x,y) with one or more regional tags. Fourteen tags (MROIs) were used based on established major gross morphology as follows: crypt apex (APEX), crypt base (SUB-CRYPT), crypt mid (MID), crypt base and mid (BASE), cross-mucosa (CM), epithelium and muscularis mucosae (EMM), epithelium and muscularis mucosae and submucosa (EMMSUB), epithelium and muscle and submucosa (ALL), muscularis externa (ME), muscularis externa and interna (MEI), muscularis interna (MI), muscularis mucosae and interna (MMI), muscle and submucosa (MSUB), and Peyer’s patch (P).

### Detection of tissue sections in histology images

In most cases, more than one tissue section was placed on the active area of one ST array. To distinguish between different tissue sections, a two-dimensional integer lattice was assumed so that labeled ST spots that were connected were assigned the same tissue section. Next, ST spots were filtered based on their sequencing data quality, such that tissue sections labeled with less than 5 (ages 0d-3w) or 10 (ages 4w-2yrs) spots in total were discarded from further analysis. ST spots with less than 800 UMIs were also discarded from further analysis. To account for spots without 4 neighbors, each spot was mapped after filtering to match the same two-dimensional integer lattice [[0,1,0],[1,1,1],[0,1,0]] and spots not matching this patterns were also discarded.

### Training a cell density classifier for segmentation

To train a cell density classifier to segment individual objects in the histology images, each whole slide image (WSI) was first subset into smaller patches while retaining patches at the same resolution for training, selected such that each patch would contain all the major colon layers from at least one tissue cross section from the original WSI. Overall, ∼200 patches were selected for training, with at least 10 replicate patches from each of the different ages. To count the number of cell segments present in each ST spot, a density classifier was first trained using Ilastik^154^. This workflow estimated the density of blob-like structures usually present as overlapping instances, decreasing the chance of underestimating the number of objects due to under-segmentation, which we reasoned was the most appropriate approach for counting cell dense areas in the colon. To ensure reproducibility across all density conditions in the dataset, in each training patch, at least three separate tissue areas (*i*.*e*. training squares) were used. In each training square, two classes of objects were labeled: cells and background.

### Processing segments per ST spot

Each WSI processed with the SpoTteR spatial transcriptomics processing tool^153^ was split into image patches of 200×220 pixels representing the size of an ST spot capture area. The cell counting workflow described above was then used to extract density predictions for each ST spot. The following image processing and segmentation steps were performed with ^155^Skimage^155^ (v.0.18.1). First, an ellipse shape (radius = 100 px) was used to mask the true ST capture area in each patch. If no cells were left in the patch (mean image intensity <0.05), the patch was discarded from further processing. Next, the multi-Otsu^156^ thresholding algorithm (cut off >50) was used to separate objects detected in the patch. Local maxima were found for each object and used to estimate distances between the same. These were then approximated by the watershed algorithm^157^ into segments that were further labeled into individual objects used in all downstream analyses.

### Training an object classifier to obtain superclass cell type labels from histology images

An object classifier was trained using Ilastik^154^, with binary segmentation images and their corresponding H&E patches as input. In this way, each segment in the H&E patch was assigned a cell type superclass label. Five different classifiers (one per cell superclass label) were created for 14 MROIs present in the colon data, separately for juvenile (<6w) and adult (>8w) groups. To train each classifier, ∼150 patches were randomly selected from all three regions of the colon and from each of the following MROIs: CM, EMMSUB, SUB-CRYPT, MID, APEX, MEI and PP. In each classifier, depending on the cells in the MROI, up to five superclasses were labeled: Colonocyte, Immune, Interstitial, Muscle and Epithelial. The object classifiers take into consideration object-level characteristics, such as object shape and work to predict similar objects in the nearby space. Algorithm features used in training included 2D convex hull and 2D skeleton descriptors in a neighborhood size of 30×30 pixels for each object, and used a simple threshold (0.5) with a small smoothing factor (1.0). Properties attributed to standard object features such as shape, size, channel intensity and location were also selected in the training process. In total, 1,540 patches and 83,721 segments were labeled during training. MROI-specific classifiers with corresponding cell type superclass labels and snRNA-seq cell type labels are presented in **Supplementary Table 3**.

### Processing cell type superclass from histology images per ST spot

Each H&E image patch (200×220 pixels) and corresponding segmentation predictions was used in Ilastik batch processing to predict cell type superclasses using the object classifier described above. Cell label predictions were used in the following image processing workflow implemented using skimage^155^ (v.0.18.1). Each pixel class in the image was assigned one of the cell type superclasses. Then, small objects (<50 px) were removed from each patch, and the remaining small segments in close proximity to each other were merged if belonging to the same cell type class. The fraction of foreground pixels belonging to each object class were used as estimates of the abundance of each cell type in each patch.

### Testing the object classifier used for obtaining superclass cell type labels from histology images

To evaluate the performance of the cell classifiers, a test set of 781 patches spanning the five adult and five juvenile classifiers was set aside. Foreground objects were detected using the binary segmentation workflow, after which all objects were manually assigned to one of the five superclasses (Colonocyte, Immune, Interstitial, Muscle and Epithelial). The same images were then input into the respective object classifiers, and confusion matrices were calculated between the manual labels and the predictions.

### Morphology-informed deconvolution using SPOTlight

The SPOTlight model was used for “bottom-up” deconvolution of ST data^48^ that takes as input two matrices of count data: *V*, a (genes x cells) matrix containing the snRNA-seq count data (in which each cell is assigned to one of *k*_*sn*_ types), and *V′*, a (genes x spots) matrix containing the ST count data. Expression matrices were pre-processed in the following manner: (1) genes were subset to the set shared across both modalities, (2) data were depth-normalized to 10k UMI counts per cell/spot and (3) data were scaled gene-wise to unit variance. Next, genes were further subset to cellular marker genes (log_2_(fold change)>1; B-H FDR <0.05; Likelihood ratio test) and balanced across the *k*_*sn*_ cell types, selecting the top *m* = *23* genes by FDR for each cell type where *m* was chosen as the minimum number of significant marker genes across all cell types (23 for T-cells). This resulted in a total of 334 unique genes used in deconvolution (**Extended Data Table 4**).

*V* was factored into component matrices *W* (genes x topics) and *H* (topics, cells) by non-negative matrix factorization (NMF):

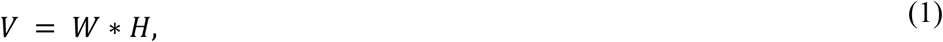

where the number of topics is assumed to be equal to the number of snRNA-seq cell types *k*_*sn*_. Prior to NMF, all *W*_*g,t*_ were initialized to the probability of gene *g* being a marker gene for cell type *t* (quantified as BH-corrected *P*_*adj*_ of *t*-test on log(count) data), and *H* was initialized as a binary matrix denoting the class assignment for each cell in the dataset. These initialization conditions – in which topics were treated equivalently to cell types – are meant to bias the optimization towards the discovery of biologically meaningful topic profiles.

Next, topic profiles *W* were fixed, and the following equation:

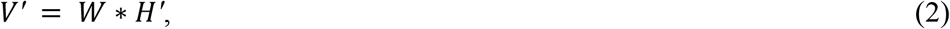

was minimized over *H′* (topics, spots) by non-negative least squares (NNLS). In this manner, the expression profile of each spot in the ST data was mapped to a combination of topics inferred from snRNA-seq.

Third, *Q*, a (topics, *k*_*sn*_) matrix from *H* was derived by selecting all cells from the same cell type and computing the median of each topic for a consensus cell-type-specific topic signature. This topic matrix was used in a final NNLS minimization to find *P*, the (*k*_*sn*_, spots) matrix denoting the inferred cellular composition of each ST spot:

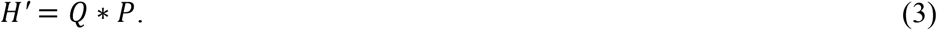

A modification to **Equation 3** was implemented that allows the incorporation of morphology-informed composition information derived from the image segmentation workflow, by providing two additional inputs: *L*, a (*k*_*morph*_, spots) matrix containing the composition of each ST spot in terms of the *k*_*morph*_ morphological cell types defined in the segmentation model, and *S*, a (*k*_*morph*_, *k*_*sn*_) binary matrix mapping each expression cell type to a morphological cell type. Any proposed compositional matrix *P* should additionally satisfy the following:

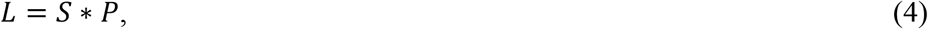

in order to reconstruct morphology-informed compositional data. Morphology-aware SPOTlight decomposition was then achieved by solving the following optimization problem:

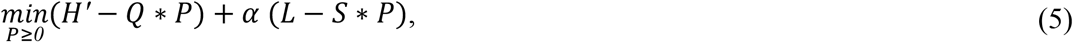

where *α* controls the relative importance of the morphological composition loss (second term) and the expression loss (first term). This optimization problem was solved using the PyTorch implementation of the Adam optimizer with a learning rate of 0.01 run for 100,000 iterations from a random initialization.

### cSplotch model specification

Genes *i*, tissue sections *j*, and spots *k* were indexed as follows: *i* ∈ [*1*, ⋯, *N*_*genes*_], *j* ∈ [*1*, ⋯, *N*_*tissues*_], *k* ∈ [*1*, ⋯, *N*^(*i*)^_*spots*_]. Each tissue *j* was registered to a common coordinate system, such that each spot *k* was assigned to one of *N*_*MROI*_ distinct MROIs, denoted 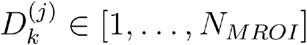, as described in the “*Spatial Transcriptomics spot annotation”* section. In the compositional mode, each spot was additionally assigned a simplex vector 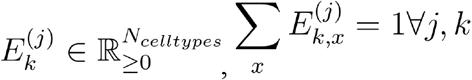, that describes its proportional composition in terms of all *N*_*celltypes*_ unique cell types across the dataset.

For each gene and each spot, the observed counts *y*_*i,j,k*_ were considered to be realizations of random variable with an expected value equal to *s*_*j,k*_*λ*_*i,j,k*_, where *s*_*j,k*_ is a size factor (total number of UMIs observed at spot *k*), and *λ*_*i,j,k*_ is the rate of expression of gene *i* (events per exposure), such that gene expression is modeled independently of sequencing depth. In practice, *s*_*j,k*_ was further normalized by the median depth across all spots in the dataset in order to facilitate comparisons of results across analyses. Thus, cSplotch offers the user two choices for modeling count data: the Poisson distribution or the negative binomial (NB) distribution. Either may be supplemented with zero-inflation to account for dropout events (technical zeros), yielding the zero-inflated Poisson (ZIP) or zero-inflated negative binomial (ZINB) distributions:

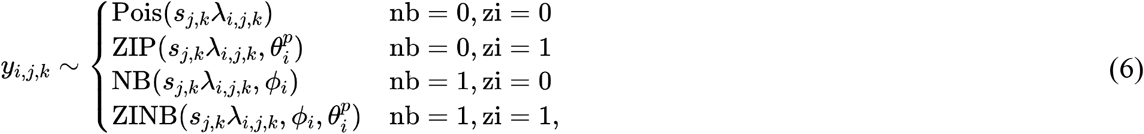

where *s*_*j,k*_*λ*_*i,j,k*_ represents the expected mean of all distributions, *ϕ*_*i*_ represents the gene-specific over-dispersion parameter for the NB family, and 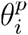 represents the gene-specific probability of technical zeros/dropout. The zero-inflated model account for an overabundance of zeros by introducing a second zero-generating process gated by a Bernoulli random variable:

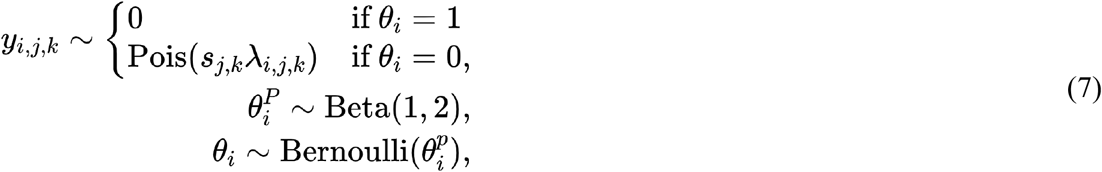

where the Poisson process can be replaced by NB without loss of generality. This mixture model allows for “true” biological zeros to be generated by the Poisson/NB process describing the expression model, while “shunting” technical zeros into a separate, technical process, preventing abundant dropout events from lowering the estimated mean expression *λ*_*i,j,k*_. Because the Poisson process does not allow for over-dispersion (variance exceeding the mean), ZIP should be preferred to Poisson in most situations, while use of NB or ZINB may depend on data quality.

While cSplotch considered a separate random variable to describe gene expression in each spot, the rate parameters *λ*_*i,j,k*_ were described in terms of a generalized linear model (GLM) that separates variation into shared and individual components. Namely, the rate of gene expression was informed by three components:

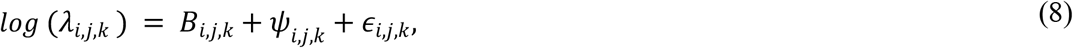

where *B*_*i,j,k*_ describes the characteristic expression of gene *i* within the tissue context of spot *k, *ψ**_*i,j,k*_ describes the neighborhood effects, and *ϵ*_*i,j,k*_ describes spot-specific effects. *B*_*i,j,k*_ is calculated as a weighted sum of cellular expression rates *β*_*i*_ in proportions *E*_*k*_^(*i*)^. Cellular expression was allowed to vary both across MROIs and sample conditions. As such, a characteristic expression `matrix` 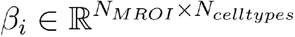 was defined (when compositional data are unavailable, each MROI may be treated as composed of a single “average” cell type and 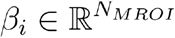 is defined instead). Inferring a posterior over *β*_*i*_ allowed quantification of expression changes across regions or cell types by comparing relevant entries.

Because characteristic expression is expected to vary across conditions (*e*.*g*., age, colon region, sex), region-specific expression *β*_*i*_ was modeled in a hierarchical fashion defined by sample covariates. Up to three levels were explicitly modeled in the hierarchy, each of which split the sample to distinct groups along some covariate. At the top level, the dataset was split along an important covariate (*e*.*g*., age), and a separate 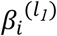 was modeled for each unique set *l*_*1*_ ∈ {*1*, ⋯ *L*_*1*_}. At the next level, each set was further partitioned along another covariate (*e*.*g*., colon region). 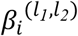 was assumed to be centered around its corresponding top-level estimate 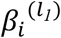, with some additional variance associated with the new covariate 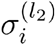. This encoded knowledge about the experimental system, and separated out sources of variation associated with each covariate. A three-level hierarchical model for *β*_*i*_ was thus specified as:

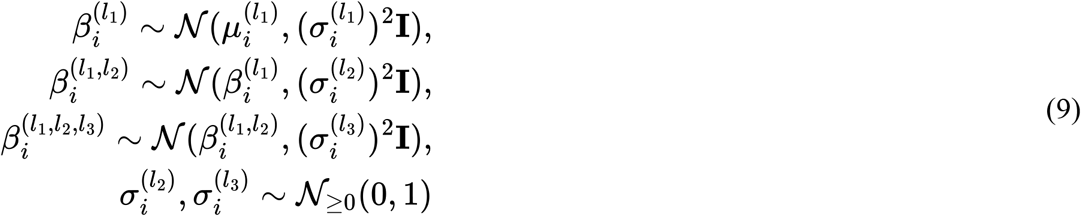

where in the compositional mode, prior hyperparameters 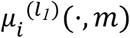 and 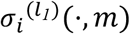 are set to the empirical mean and standard deviation (respectively) over the expression of gene *i* in cell type *m* in the snRNA-seq data for all MROIs, and in the non-compositional mode (one “average” cell type per MROI) 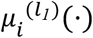 and 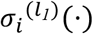 are set to 0 and 2, respectively, for all MROIs. Variation parameters 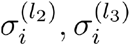 are assumed to have truncated Gaussian priors reflecting our limited knowledge of the effects of covariate-driven variation, and inferred separately for each level 2 and 3 covariate group. For convenience, because each tissue *j* belongs to one covariate group at each level, the inverse mapping function *ρ*^−*1*^(*j*) was introduced that maps *j* to the appropriate *l*_*1*_, *l*_*2*_, *l*_*3*_ indices for *β*_*i*_. *B*_*i,j,k*_ was formally defined in the non-compositional model:

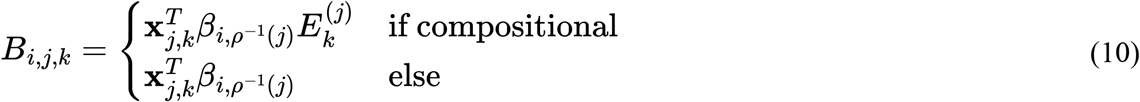

Where 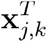 is a one-hot encoding of *D*_*k*_^(*i*)^, the MROI annotation for spot *k*. With this framework for integrating multiple sections or experiments the posterior distributions of the latent parameters *β*_*i*_ were studied at different levels of the hierarchical experimental design, and expression changes were quantified across conditions, tissue contexts, or individual cell types.

The second component of Equation 8, *ψ*_*i,j,k*_, describes the effects of the local neighborhood of spot *k* on the expression of gene *i*. This was modeled using the conditional autoregressive (CAR) prior, which assumes that the value at a given location (spot) is conditional on the values of neighboring locations (spots). *ψ*_*i,j*_ was defined as a Markov random field over the spots on each array:

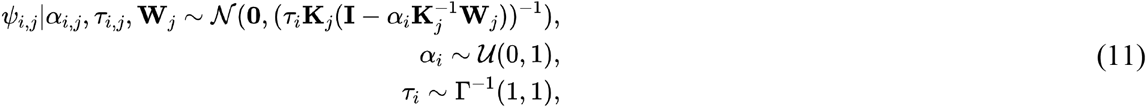

where *a*_*i*_ is a spatial autocorrelation parameter, *τ*_*i*_ is a conditional precision parameter, **K**_*j*_ is a diagonal matrix containing the number of neighbors for each spot in tissue *j*, and **W**_*j*_ is the adjacency matrix (with zero diagonal). If the classic ST methodology of utilizing cartesian arrays is employed, each spot is assumed to have a 4-spot neighborhood, while if the Visium platform utilizing hexagonal arrays is employed, each spot is assumed to have a 6-spot neighborhood. The level of spatial autocorrelation (*a*_*i*_) and conditional precision (*τ*_*i*_) was inferred separately for each gene. Taken together, the *B* and *ψ* terms capture spatial autocorrelation on two different scales: tissue context (across samples) and local neighborhood (within samples).

The final component of Equation 8, *ϵ*_*i,j,k*_, captures variation at the level of individual spots. This variation was assumed to be independent and identically distributed (i.i.d.) for each gene:

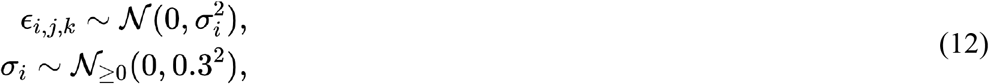

where *σ*_*i*_ is the inferred level of variability for gene *i*.

### Bayes factor estimation for cSplotch differential expression analysis

To quantify difference in expression between two conditions using cSplotch, the Bayes factor between posterior distributions over characteristic expression coefficients *β* estimated by the model was examined. Without loss of generality, difference in expression was quantified between conditions represented by *β*^(1)^ and *β*^(2)^, which may differ across any combination of genes, sample covariates (*e*.*g*., distal *vs*. proximal colon), tissue regions (*e*.*g*., crypt apex *vs*. muscle), or cell types (*e*.*g*., neuron *vs*. myocyte). A random variable **Δ**_*β*_ = *β*^(1)^ − *β*^(2)^ was defined, which captures the difference between *β*^(1)^ and *β*^(2)^. If **Δ**_*β*_ is tightly centered around zero, then the two distributions are very similar to each other, and the null hypothesis of identical expression cannot be rejected. To quantify this similarity, the posterior distribution **Δ**_*β*_|**𝒟** (where **𝒟** represents the data used to train the model) was compared to the prior distribution **Δ**_*β*_ using the Savage-Dickey density ratio^158^ that estimates the Bayes factor between the conditions:

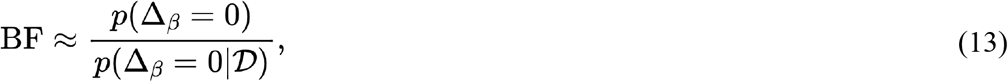

where the probability density functions were evaluated at zero. If expression is different between the two conditions, then the posterior **Δ**_*β*_|𝒟 will have very little mass at 0, and the estimated Bayes factor will be large (by convention, BF > 5 indicates substantial support). Conversely, for similar expression regimes, the posterior will place a mass equal to or greater than that of the prior at zero, and the Bayes factor will be ≤ 1. While *p*(**Δ**_*β*_) can be derived analytically (the prior distributions over all *β*_*s*_ are normally distributed, and the difference between two normally distributed random variables is in turn normally distributed), *p*(**Δ**_*β*_|**𝒟)** must be approximated using the posterior samples obtained in the following section. When we executed a comparison between sets of conditions (e.g., neurons vs. all other cells), we pooled the posterior samples from all component conditions together.

### Parameter inference for cSplotch

cSplotch was implemented in Stan^159^. For all analyses, Bayesian inference was performed over the parameters using Stan’s adaptive Hamiltonian Monte-Carlo (HMC) sampler with default parameters. Four independent chains were sampled, each with 250 warm-up iterations and 250 sampling iterations, and convergence was monitored using the R-hat statistic.

### Simulated ST data generation

Simulated ST data were generated from the snRNA-seq profiles in the two regimes described in the subsequent sections. For all simulation studies, 12 ST arrays were generated, each containing 2,000 spots. For data in which distinct MROIs were simulated, the two regions were considered to exist in a 1:1 ratio. Cell clusters comprising each region are detailed in **Extended Data Table 2**.

### Cluster-based simulation

Average expression profiles for unique mouse colon cell types obtained from snRNA-seq 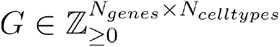 (*N*_*genes*_ = *22,986, N*_*celltypes*_ = *30*) were normalized column-wise, such that the total expression within each cluster summed to 1. For each spot *k* from tissue *j*, a “true” composition vector 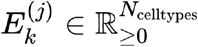 was drawn such that the cell types present in the current region 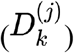 were represented in uniformly random proportions. For each cell type *t*, reads *y*_*k,t*_ were drawn from a multinomial distribution:

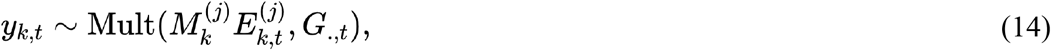

Where 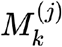 is the total number of reads in the current spot and *G*_.,*t*_ is the expression profile for cell type *t*. In practice, *M*_*k*_^(*j*)^ = *1,000* was used for all *j,k*. Read counts were then pooled across all cell types, yielding spot-level reads *y*_*k*_ = Σ_*t*_ *y*_*k,t*_. Composition vectors were then (optionally) pooled within high-level annotation categories (such as the histological superclasses in **Extended Data Table 2**) in order to simulate cases of limited observability. Finally, Gaussian noise was (optionally) added to 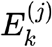 in order to generate “observed” composition vectors 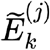. Resulting negative entries were removed by element-wise maximization against the zero vector 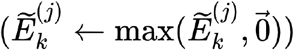, following which 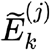 was re-normalized to produce a valid simplex. *y, D*, and 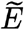 served as inputs to cSplotch.

For all cluster-based data, cSplotch was run with a Poisson likelihood (without zero-inflation) to meet the expected form of the marginals over the generating distribution (multinomial). In order to focus on the compositional module of cSplotch, the effects of local spatial autocorrelation were removed by simulating each spot independently, and as such suppressed the *ψ*_*i*_ term of the GLM. Local neighborhood effects can be simulated in this regimen by passing a spatial smoothing filter over the simulated array, blending the transcriptome of each spot with those of its neighbors in a defined proportion.

### Cell-based simulation

Individual snRNA-seq cell profiles 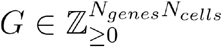 (*N*_*genes*_ = *22,986, N*_*cells*_ = *419,334*), where each cell was assigned to one of *N*_*celltypes*_ = *30* cell types, were normalized cell-wise to a target depth of 1,000 counts using Scanpy’s ^150^ `normalize_total` function. Each spot *k* on tissue *j* was comprised of 10 cells, partitioned uniformly at random among the cell types present within the current region 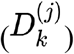. The integer vector containing the number of cells belonging to each type was normalized to produce a simplex serving as the “true” composition vector 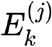. For each cell type *t* 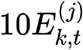, cells were drawn at random (without replacement) from *G*^(*t*)^, the subset of cell profiles that have been annotated as type *t*. Their expression profiles were summed to yield *y*_*k,y*_, then rounded to integer values to remove fractional reads introduced by `normalize_total`. As with the cluster-based approach, read counts were then pooled across all cell types, yielding spot-level reads *yk* =Σ_*t*_ *y*_*k,t*_, and composition vectors were then (optionally) pooled within high-level annotation categories. Observational noise was added in an identical fashion to the cluster-based approach. For all cell-based data, cSplotch was run with a negative binomial likelihood (without zero-inflation) in order to account for overdispersion present over gene counts in the snRNA-seq profiles. As with the cluster-based approach, spots were assumed to be fully independent and thus *ψ*_*i*_ was suppressed in the GLM.

### Power sampling and its effect on estimation

ST data of distal colon sections from 12w-old mice were chosen for sub-sampling analysis given the large number of samples from these mice (six mice (three males; three females) with 13, 8, 9, 8, 8, and 6 tissue sections per mouse). To sub-sample the data, a given number of mice were randomly selected, and a given number of tissue sections selected from each mouse (if a selected mouse had less than the given number of tissue sections, all tissue sections belonging to that mouse were taken). Each sub-sampled data set (nine combinations of 1, 2, and 6 mice with 2, 4, and 8 tissue sections per mouse) and the full data set (52 tissue sections and six mice) were analyzed separately by cSplotch. To compare the estimated posterior distributions of *β*_*i*_ at the gene and tissue context levels, the posterior means and their standard deviations were calculated, which were then used to obtain normal distribution approximations of the posterior distributions. The Kullback-Leibler divergence (KLD) between the posteriors derived from the full and sub-sampled data (*i*.*e*., how much information is lost when using the distribution estimated from the sub-sampled data) was used to quantify the differences between the normal distributions.

### Characterizing multi-cellular programs of gene expression

The cSplotch model output of posterior mean estimates of cellular gene expression 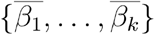 for *k* cell types in a given spatial niche (*e*.*g*., the *crypt apex* MROI), where each 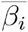 is a matrix of dimension *N*_*genes*_ × *N*_*conditions*_ was used as input to identify gene sets spanning multiple cell types that show *k* -way correlation across the measured conditions. First, for each spatial niche, cell types and genes were selected for MCP analysis. Only cell types that are found in at least 5% of spatial spots, on average, were included, in order to focus on cell types with sufficient certainty in posterior estimates of gene expression. Across the *k* included cell types, the top 5% of gene signatures were considered based on coefficient of variation of *<u>β</u>* across the conditions, to focus on genes with spatio-temporal variation.

Next, penalized matrix decomposition (PMD) was used to find linear combinations of gene signatures in each cell type that are maximally correlated across all conditions^160^. PMD seeks to find sparse canonical variates {*w*_1_, …, *w*_*k*_} that transform the original gene feature space into a new “MCP” feature space of dimension *M* ≤ *k* such that:

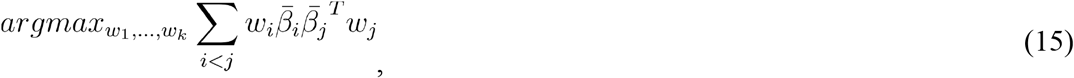

subject to ∀_*i*_‖*w*_*i*_‖ ≤ 1, *ρ*_*i*_ (*w*_*i*_) < *c*_*i*_, where *ρ*_*i*_(*w*_*i*_) represent LASSO regression penalties, and *c*_*i*_ controls the degree of sparsity. These tuning parameters were chosen by a permutation-based approach as previously described^54^. For each pair of cell types *i* and *j*,

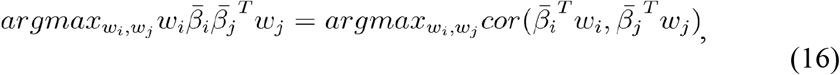

therefore the canonical covariates identified a space in which each MCP signal in cell type *i* is highly correlated with the corresponding signal in all other cell types. Next, given canonical variates from PMD, each of the *M* MCP signals was characterized by identifying at most n=250 genes across all cell types that show the strongest positive or negative contribution as measured by *abs*(*w*) > *0*.*1*. The list of correlated genes for each MCP – both “up” genes with positive weight and “down” genes with negative weight – was then used as input to a Fisher’s exact test to identify KEGG functional gene sets enriched in the MCP. Each MROI was allowed at most *k* MCPs under the constraint that the bases for each MCP are orthogonal. As MCCA will always output programs in the same order, the maximum of *k* MCPs was calculated for a given spatial niche and latter programs that show low self-correlation among member genes (mean Pearson r < 0.3) or high cross-correlation with genes from other programs (mean Pearson r > 0.05) we optionally removed. Total MCP activity across conditions was calculated as a weighted sum of member gene activity 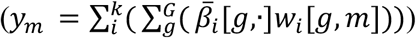, while cellular contributions to each MCP were calculated by binning positive and negative weights by cellular origin.

### Spatio-temporal cellular composition analysis

To identify significant differences in cellular composition, each tissue section was treated as an independent sample, and compositional estimates of all spots for a given MROI were pooled from the same section. Statistical significance across groups (*e*.*g*., proximal-middle-distal samples) was assessed using Welch’s t-test. A crypt gradient gene was defined as one that showed a significant (BF > 2, LFC > 0.5) difference in expression between crypt base and apex, and a monotonic change in expression from base to base & mid, mid, and apex.

## Data availability

All data have been deposited in the Single Cell Portal under accession SCP2595 (https://singlecell.broadinstitute.org/single_cell/study/SCP2595).

## Code availability

All code is deposited on GitHub (https://github.com/adaly/cSplotch). A Google Cloud-enabled workflow is also available on Terra Firecloud (https://app.terra.bio/#workspaces/techinno/cSplotch%20Workflow)

## Acknowledgements

Work was supported by the Knut and Alice Wallenberg Foundation, Beijer Laboratory for Gene and Neuro Research, the Royal Swedish Academy of Sciences, Swedish Society for Medical Research, Science for Life Laboratory, 1U54 AG076040-01 and 1RM1 HG011014-01 (S.V.), and the Klarman Cell Observatory, the Manton Foundation, and HHMI (A.R.). S.V was supported as a Wallenberg Fellow at the Broad Institute of MIT and Harvard and as a Wallenberg Academy Fellow and SciLifeLab Fellow at Uppsala University. A.R. was an Investigator of the Howard Hughes Medical Institute. We would like to thank the Flatiron Institute for providing computing resources that enabled the completion of this work.

## Author contributions

S.V. conceived and designed the study with guidance from A.R. and R.B.; S.V. performed the experiments with help from B.L., N.D.M., S.F., N.v.W. and E.D.; A.D. analyzed the data with guidance from S.V. and R.B. and help from T.Ä., M.S-E and B.L.; S.V. annotated the histological sections with help from O.K. and N.v.W. with guidance from G.G.; S.V., A.D., F.C. and A.R. interpreted the data and wrote the manuscript with input from all the authors. All authors discussed the results.

## Extended Data Figure legends

**Extended Data Fig. 1: Cell type associated genes from snRNA-seq**. Mean expression (dot size) and fraction of expression cells (dot color) of the top three marker genes (columns) for each of the 17 cell subsets (columns) defined from snRNA-seq.

**Extended Data Fig. 2: Validation of semantic segmentation of morphological cell types (“super-classes”)**. (**a,b**) Semantic segmentation workflow. Semantic segmentation of cell nuclei (**c**), conditioned on five broad annotation groups for spots (**b**). (**c,d**) Accuracy of cell type composition by semantic segmentation. Fractions of FG pixels in each H&E image (y axis) assigned to each morphological superclass (x axis) in test and train sets (color code) in adult (c, top) and young (d, top) mice, and percent and number (color scale) of pixels with a ground truth superclass label (rows) classified to each label (columns) in in adult (c, bottom) and young (d, bottom) mice. In Box plots: center black line, median; color-coded box, interquartile range; error bars, 1.5x interquartile range; black dots; outliers. Numbers of images used in train and test sets are denoted by “n_train” and “n_test”, respectively, above each subplot.

**Extended Data Figure 3: Effect of morphological constraints on count-based deconvolution of ST data with NMF**. (**a**) Reconstruction of expression by NMF^48^ is unaffected by constrained weight parameter (*α*). Distribution (y axis) of mean squared error (MSE, x axis) between observed (N=11761 genes) and SPOTlight-reconstructed expression (**Methods**) for N=69,721 spots at different values of *α* (color code). **(b)** Cell compositions proposed by NMF decrease in entropy with increased *α*. Distribution (y axis) of entropy of cell composition vectors predicted by constrained NMF as in **(a)**. Entropy (x axis) is calculated across all 17 snRNA-seq cell types (*k*_*sn*_ = *17*), and thus ranges between 0 (one cell type present) and 1 (all cell types present in equal proportion). (**c**) Reconstruction of morphological cell compositions is greatly enhanced with increased *α*. Distribution of residual MSE between predicted and observed morphological cell type (“superclass”) composition vectors from SPOTlight as in (**a**). Morphological cell type vectors are calculated from the output of SPOTlight by pooling snRNA-seq cell types from the same morphological class (*k*_*morph*_, as in **Extended Data Table 2**), reducing the space from 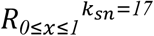 to 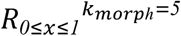.

**Extended Data Figure 4. cSplotch statistical model for ST data. (a,b)** Underlying statistical model of spatial expression for a gene *i* at spot *k* on tissue *j*. (**a**) Characteristic expression rate *β* (red shaded area) for gene *i* is inferred separately in each cell type and MROI (white rectangles). Stacked gray boxes: three-level hierarchical formulation of *β* (*l*_1_, *l*_2_, *l*_3_), where each level inherits from the one before it. Random variables ψ (yellow shaded area) and ε (blue shaded area) account for spatial autocorrelation and spot-level variation components, respectively. Gray and white circles: observed and latent variables, respectively. (**b**) Distributions over random variables of the statistical model in (**a**). ZIP, NB, ZINB: zero-inflated Poisson, binomial, and zero-inflated negative binomial distributions, respectively. (**c)** Model inputs. A multi-tissue colon dataset (left) is annotated at the spot level with MROI tags (left color key) and cellular compositions (segmentation masks, pie charts; lower color key), encoded as observed variables *D*_*k*_^(*j*)^ and *E*_*k*_^(*j*)^, respectively. The expression measurement at spot *k* (“Gene Counts”) yields the total sequencing depth of spot *k* (*s*_*j,k*_), and the observed counts (*y*_*i,j,k*_) of gene *i* (*e*.*g*., Abca8a) therein. The (4-)neighborhood of each spot on tissue *j* (right) is encoded in the adjacency matrix *W*_*j*_.

**Extended Data Figure 5. Validation of cSplotch deconvolutional capabilities on simulated ST data**. (**a,b**) Correlations between true and predicted counts for simulated cell type clusters. Recovered (y axis, 10^6^ *exp* 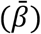, where 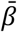 is the posterior mean expression in a given cell type) and observed (x axis, TPM, snRNA-Seq data) expression for each of 1,000 genes (dots) with real input (top row) or at different levels of corruption of cellular composition input (other three rows) in cells from each of five superclasses (columns) from a single tissue region (**a**) or in each of two cell types from each of two different tissue regions (**b**). Bottom left: Pearson’s r. Red line: *x*=*y*. (**c**) Differential expression effect size (log fold change; x axis) and significance (-log10 BH-adjusted p-value; t-test; y axis) for each gene (dot) between two cell types in one MROI (as labeled on top) based on snRNA-Seq data, with dots colored by log10 Bayes factor (BF) of an analogous DE analysis between deconvolved cell profiles from cSplotch on simulated ST. Dashed vertical lines: LFC = |1|; dashed horizontal lines: p_adj_=0.05.

**Extended Data Figure 6. Effect of experimental design on study power**. (**a**) Characteristic expression rates (*β*) in the cross-mucosa of the distal colon for 12 week-old mice. Distribution of Kullback-Leibler divergence (KLD, **Methods**) between posterior distributions of expression rates for n=12,976 genes estimated from sub-sampled data (1, 2 and 4 mice; 2, 4 and 8 tissue sections per mouse) *vs*. the full data (6 mice, 53 tissue sections). Lower KLD values indicate greater agreement between full and subsampled data. **(b**) Mean expression 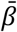 (posterior mean estimated from the full dataset) (x axis) and KL divergence (y axis) for each gene between the estimate from a sub-sampling of mice (columns) and tissue sections (rows) vs. the full data for each of n = 12,976 genes.

**Extended Data Figure 7. Effect of colon region on inferred cell-type specific expression**. Posterior distribution of characteristic expression rate (*β*; x-axis) of individual genes (labeled on top right) in each of the three regions (color) in goblet cells (left) and ISCs (right) in CM (**a**), and in SMCs (left) and fibroblasts (right) in MEI (**b**). Bold: Canonical markers. Brackets: significant differential expression (*: BF>2; **: BF>10; ***: BF>30; ****: BF>100).

**Extended Data Figure 8: Enhanced cell marker specificity after integration of snRNA-seq and ST**. Distribution of characteristic expression rate (*β*; x-axis) for canonical marker genes of select cell types (color; *Clca1, Prdx6* (goblet cells); *Pdgfra* (fibroblasts), and *Kcnq1*(TA cells)) in specific MROIs (x-axis label) from proximal (left), middle (middle) and distal (right) colon segments from 12 week old mice based on snRNA-seq data only (dashed lines; empirical prior), or as inferred by cSplotch from ST and snRNA-seq data (solid lines; posterior distributions). Only cell types present at least at 2% in MROI-annotated ST spots are included.

**Extended Data Figure 9: Spatio-temporal variation in inferred cellular composition in the crypt**. Fraction of cells (y axis; mean and standard deviation) of each abundant (>5% of spots of average) cell type (rows) at each time point (x axis) in each of four crypt MROIs (columns) in each colon region (color code). Gray area: aging window; brackets: significant changes between 12w and 2yr time points; *: 0.01 < FDR <= 0.05; **: 10^−3^ < FDR <= 0.01; ***: 10^−4^ < FDR <= 10^−3^; ****: FDR <= 10^−4^ (Welch’s *t*-test). Significant changes are also listed in **Extended Data Table 9**.

## Extended Data Tables

**Extended Data Table 1**. Composition of snRNA-seq dataset by cell type.

**Extended Data Table 2**. Composition of ST dataset by covariate group & spot annotation.

**Extended Data Table 3**. Semantic segmentation model and morphological superclass specifications.

**Extended Data Table 4**. Cell type marker genes for deconvolution.

**Extended Data Table 5**. Simulated ST data specifications.

**Extended Data Table 6**. Regional variation in mean cell composition.

**Extended Data Table 7**. Crypt gradient genes.

**Extended Data Table 8**. Cellular associations of crypt gradient genes.

**Extended Data Table 9**. Significant variations in cell composition across time.

**Extended Data Table 10**. Manually-curated functional annotation of selected MCPs along the vertical crypt axis during colon aging.

**Extended Data Table 11**. Significant associations between crypt MCPs and KEGG pathways.

