## Extended Data Figures 1-9 for "Tissue and cellular spatiotemporal dynamics in colon aging"

### Extended Figure 1

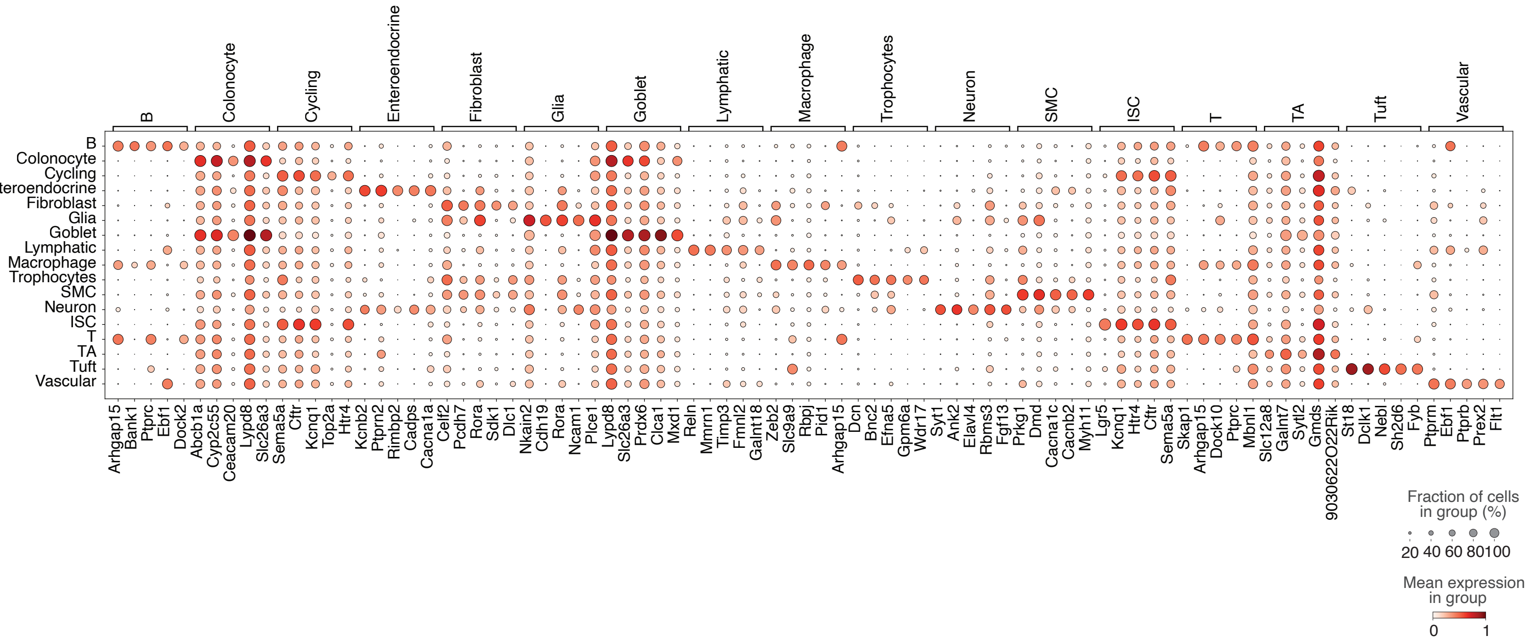

### Extended Figure 2

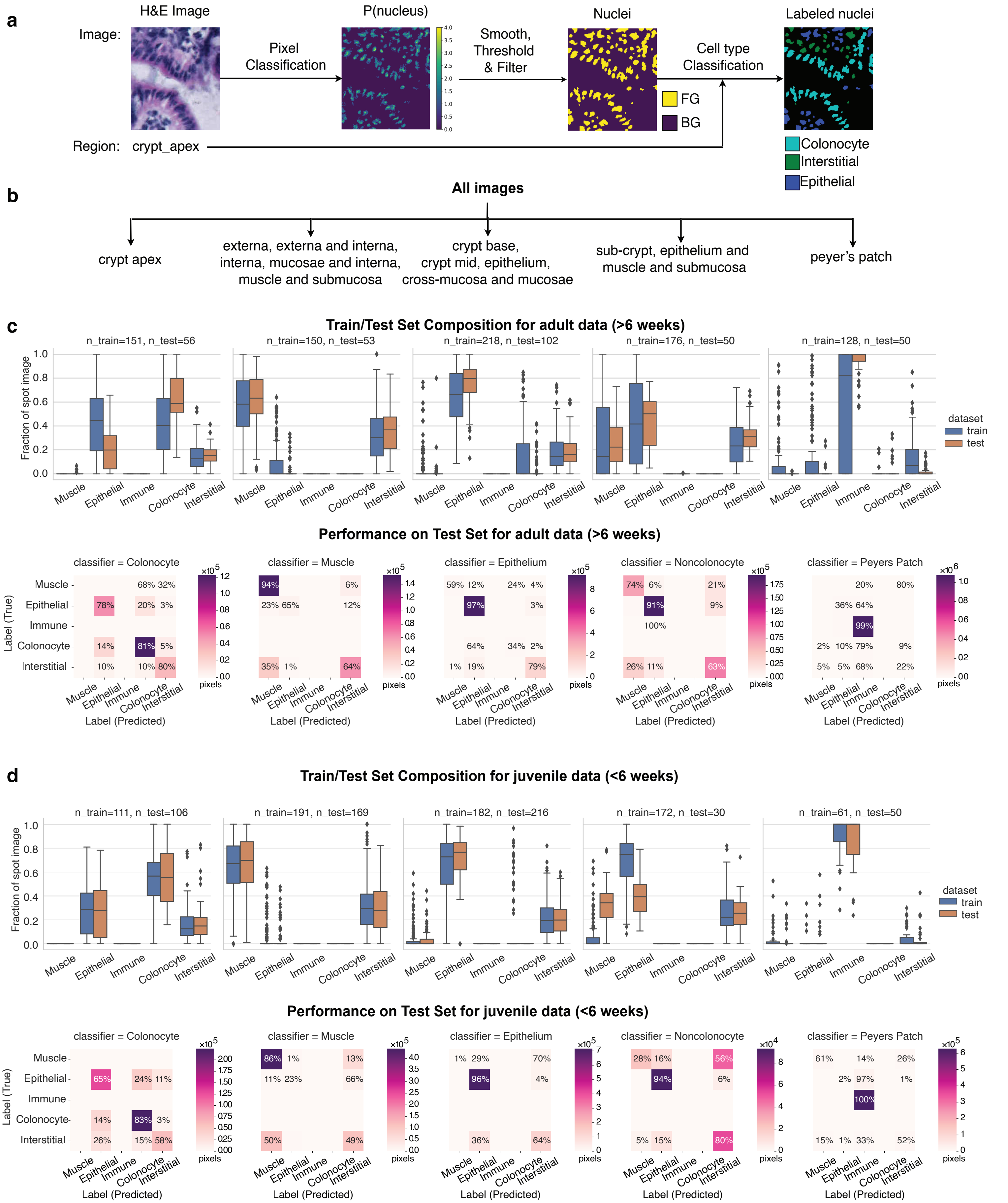

### Extended Figure 3

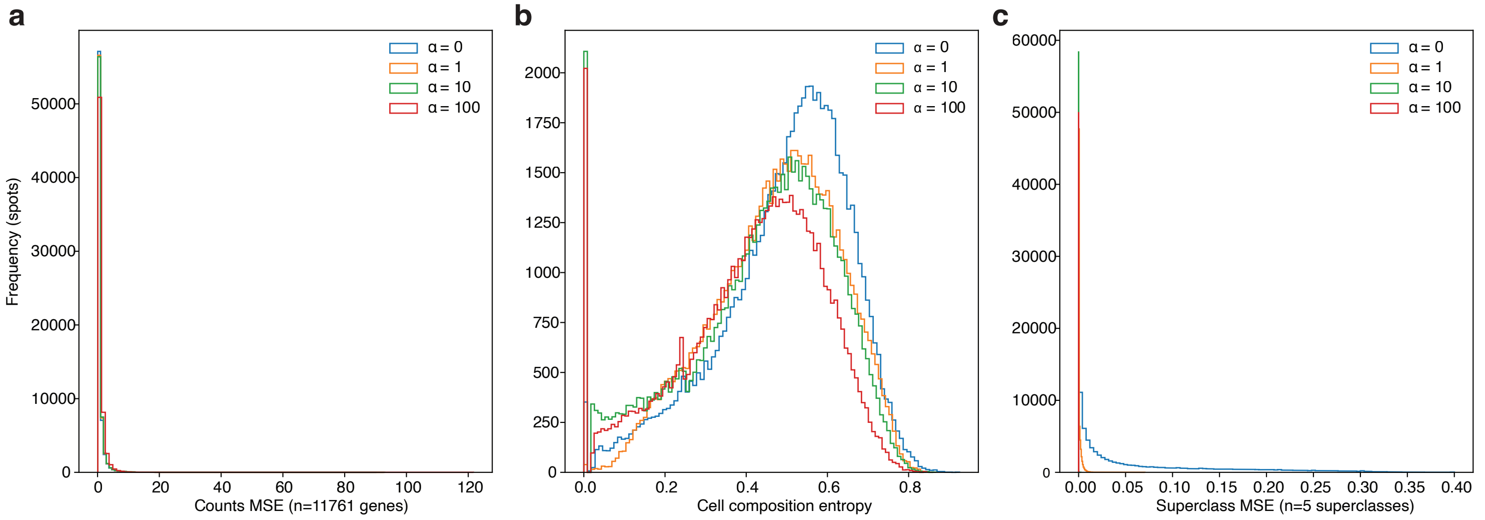

### Extended Figure 4

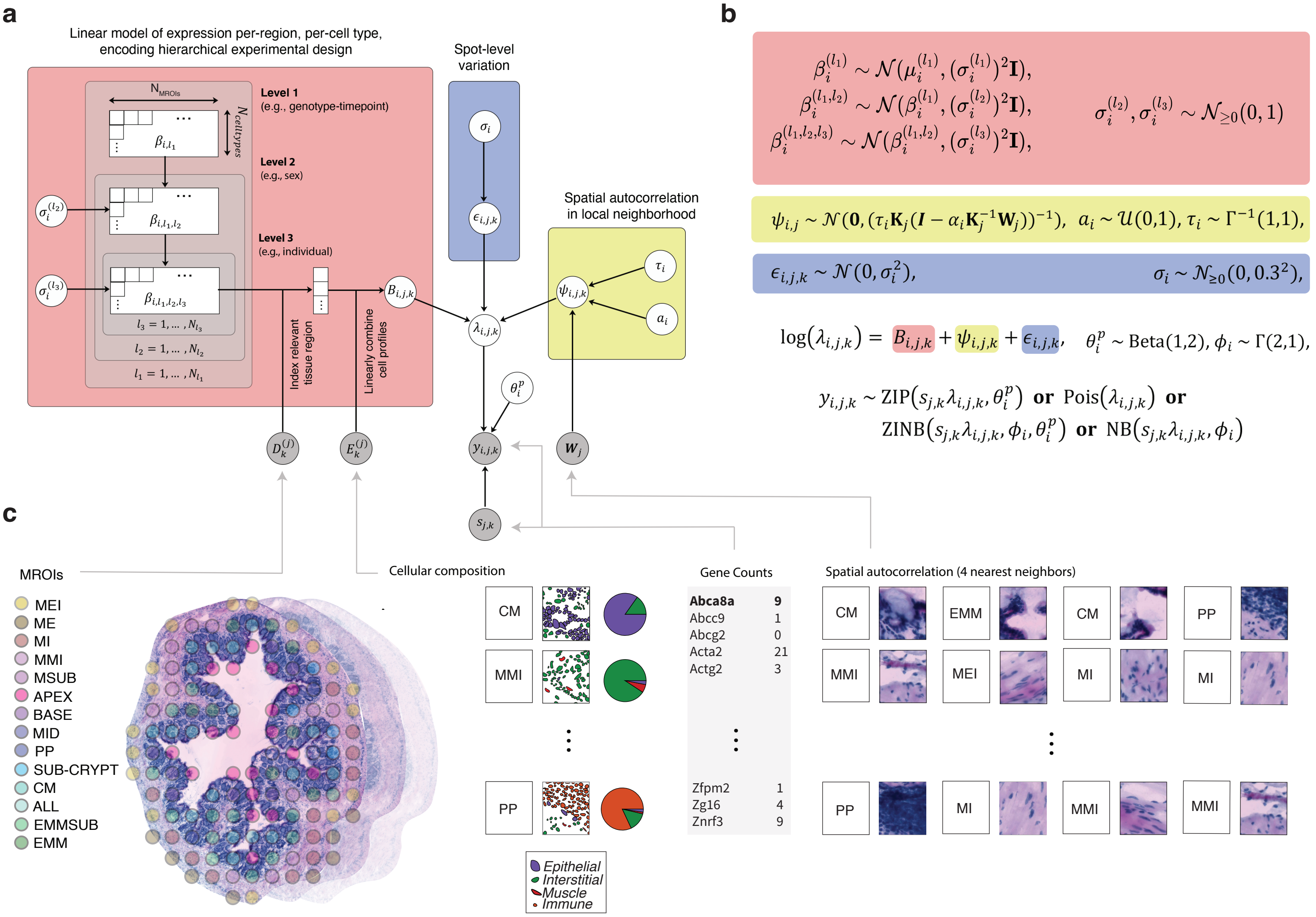

### Extended Figure 5

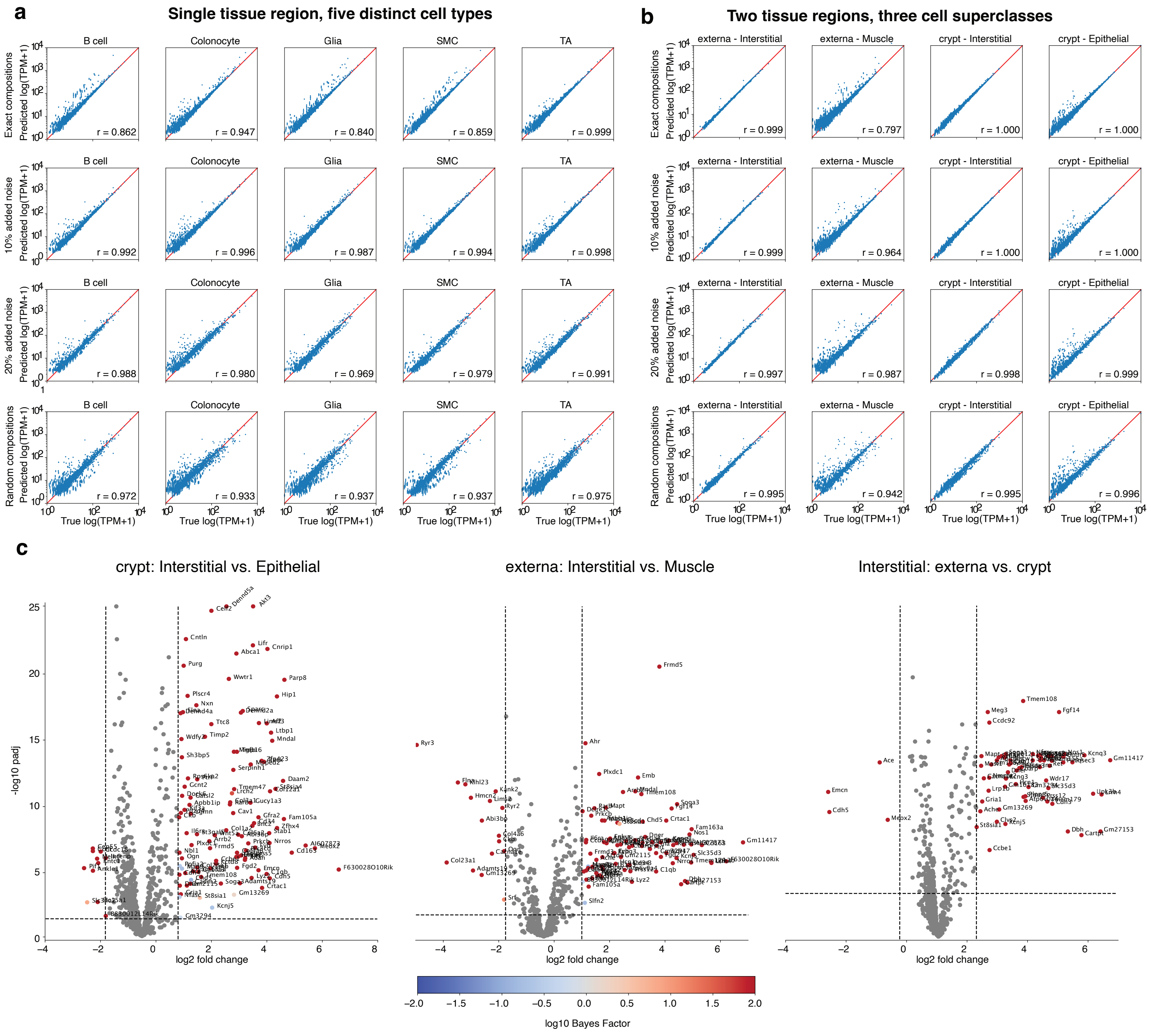

### Extended Figure 6

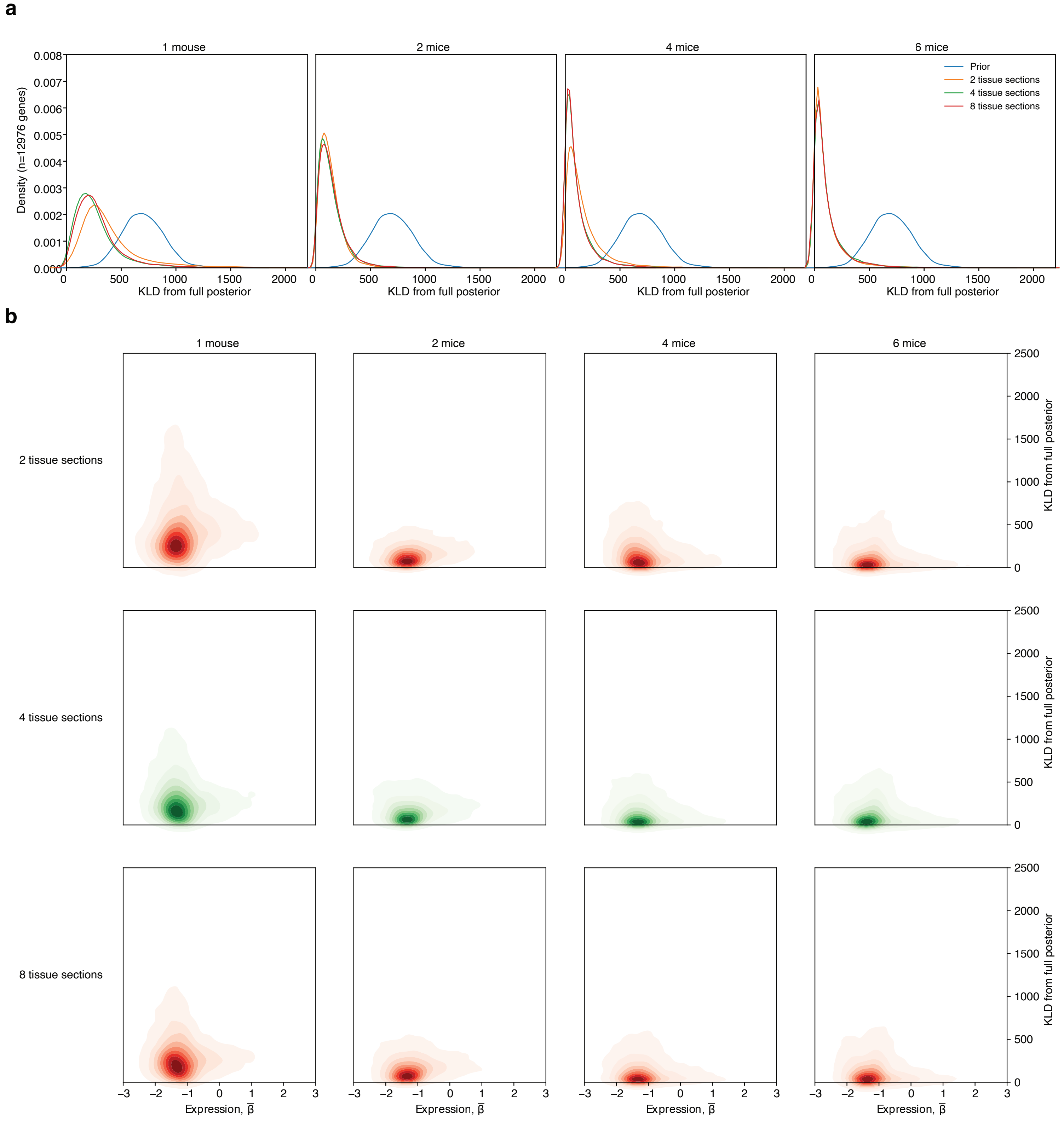

### Extended Figure 7

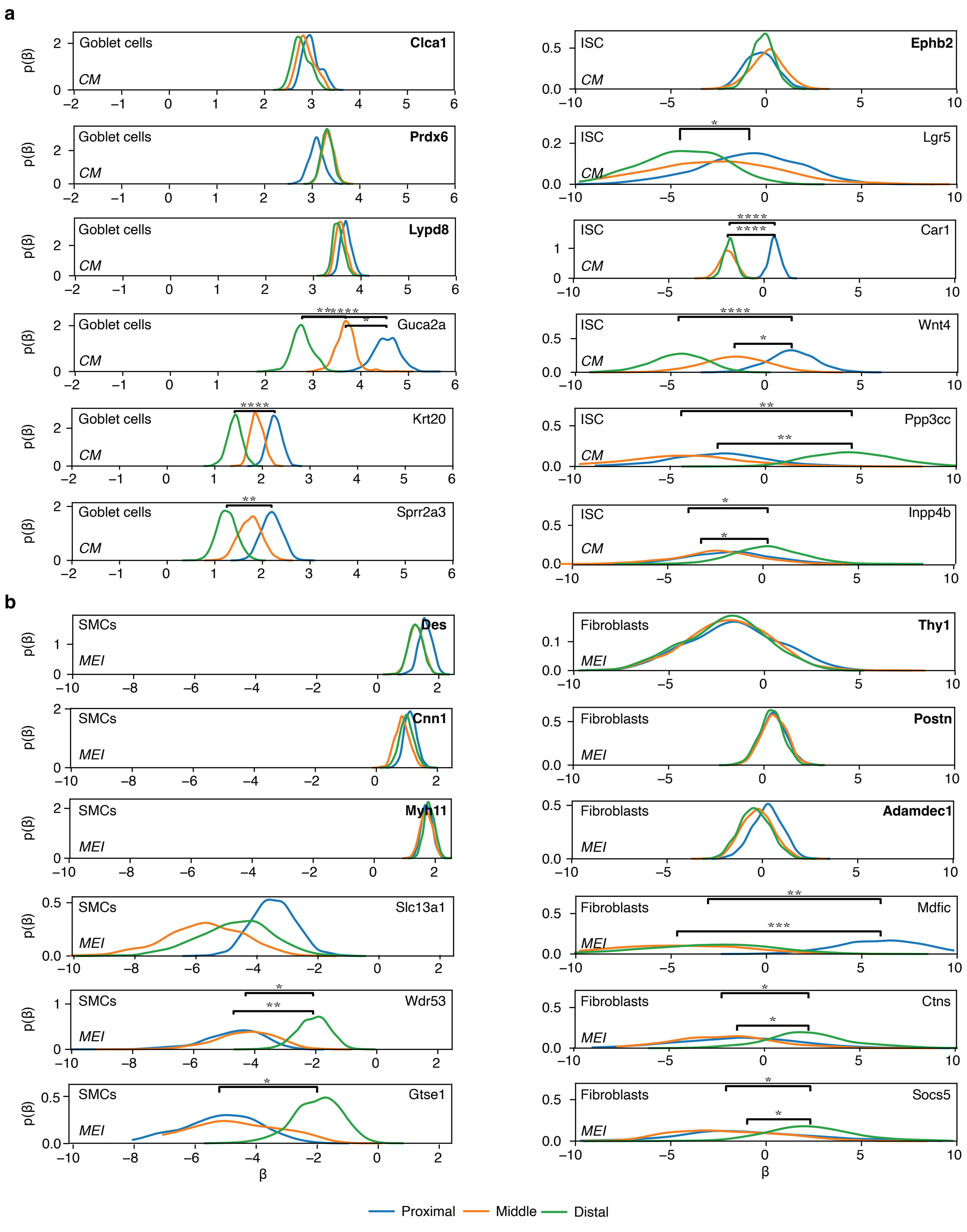

### Extended Figure 8

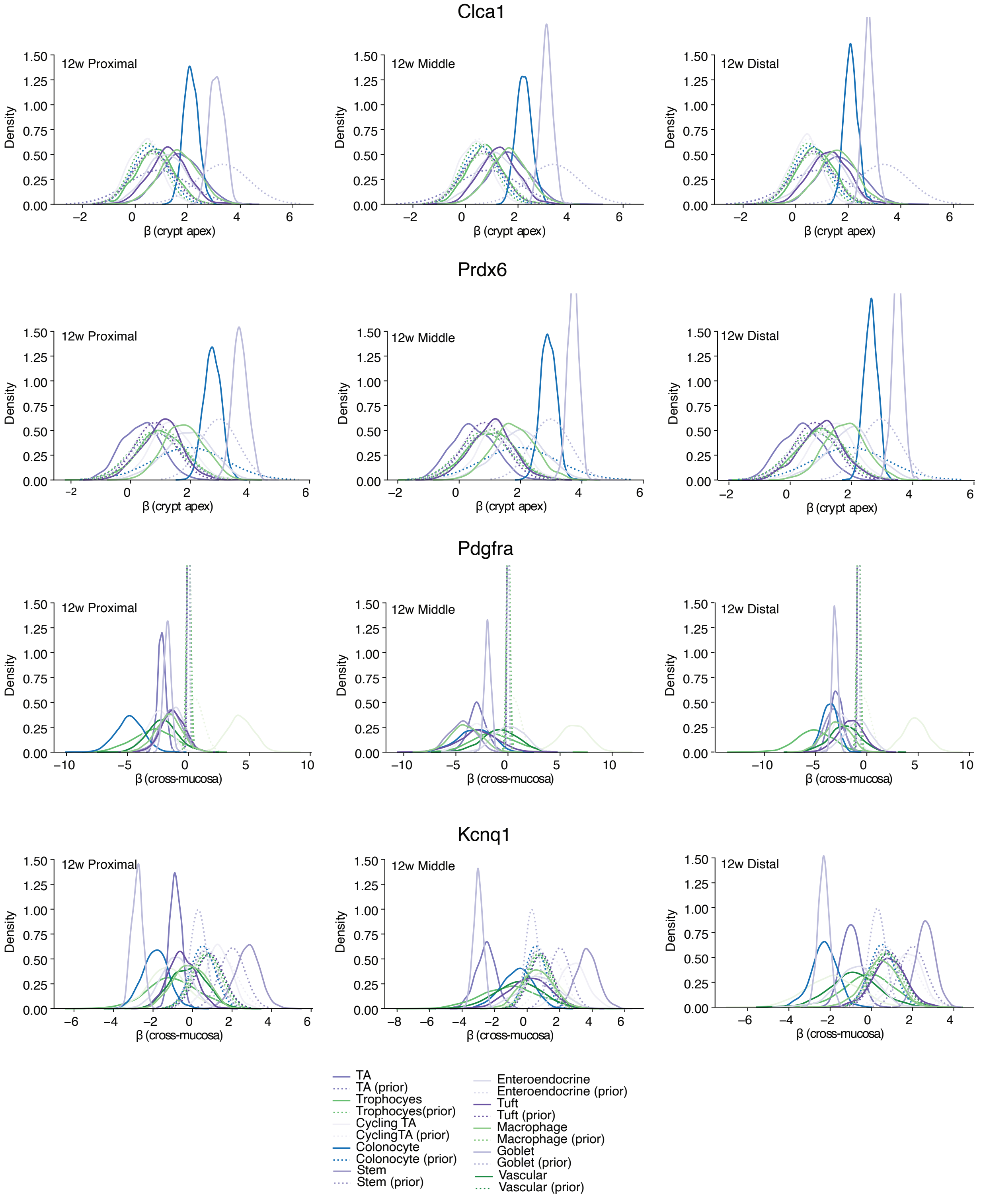

### Extended Figure 9

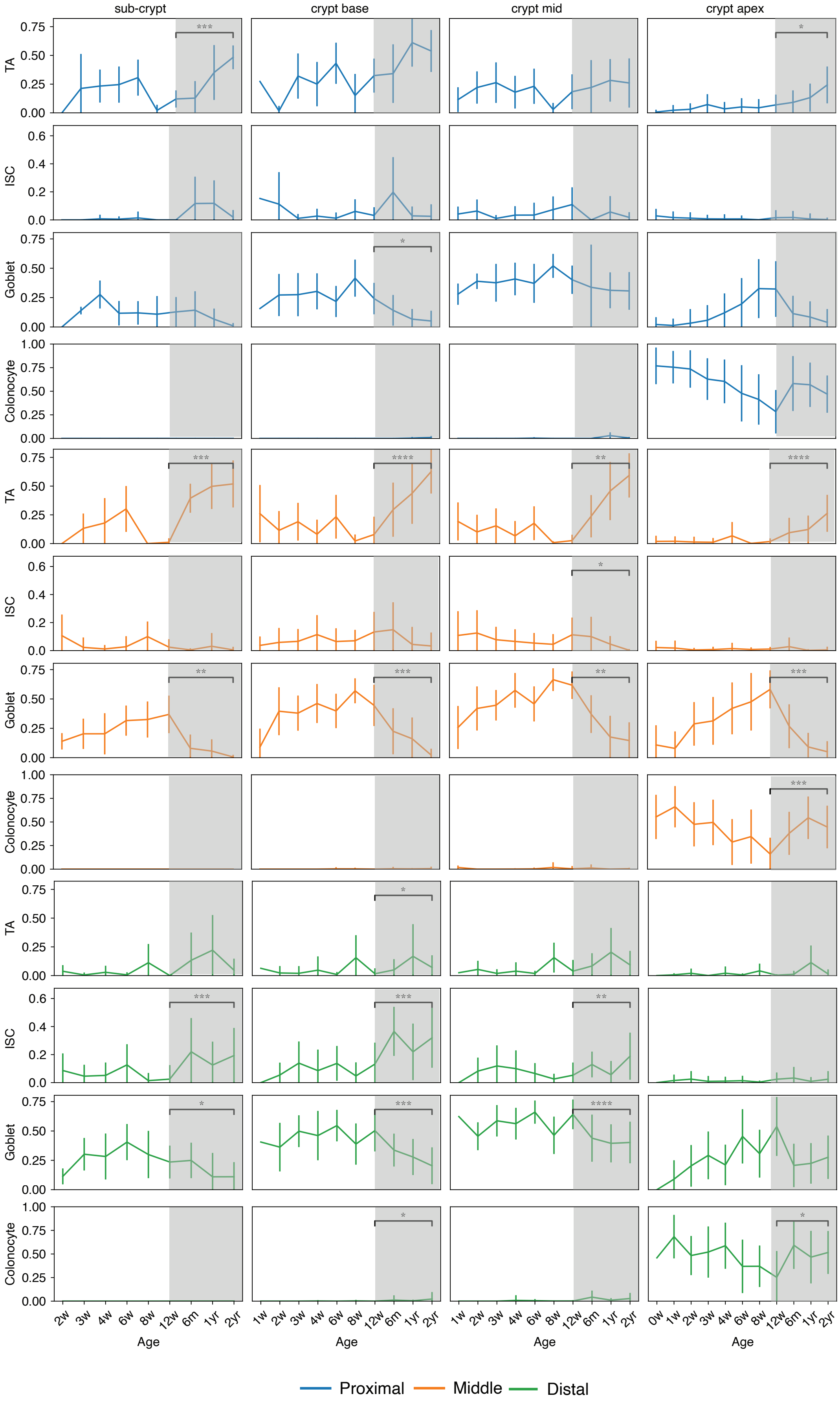
